# A Hypothalamic Circuit that Modulates Feeding and Parenting Behaviors

**DOI:** 10.1101/2024.07.22.604437

**Authors:** Ivan C. Alcantara, Chia Li, Laura E. Mickelsen, Christopher M. Mazzone, Isabel de Araujo Salgado, Claire Gao, Brian N. Papas, Cuiying Xiao, Eva O. Karolczak, Abigail I. Goldschmidt, Shakira Rodriguez Gonzalez, Ramón A. Piñol, Jian-Liang Li, Guohong Cui, Marc L. Reitman, Michael J. Krashes

**Affiliations:** Diabetes, Endocrinology, and Obesity Branch, National Institute of Diabetes and Digestive and Kidney Diseases, National Institutes of Health, Bethesda, MD, USA 20892; National Institute of Environmental Health Sciences, National Institutes of Health, Durham, NC, USA 27709; Biostatistics and Computational Biology Branch, National Institute of Environmental Health Sciences, Durham, NC, USA 27709; Department of Neuroscience, Brown University, Providence, RI, USA 20912

## Abstract

Across mammalian species, new mothers undergo considerable behavioral changes to nurture their offspring and meet the caloric demands of milk production^1-5^. While many neural circuits underlying feeding and parenting behaviors are well characterized^6-9^, it is unclear how these different circuits interact and adapt during lactation. Here, we characterized the transcriptomic changes in the arcuate nucleus (ARC) and the medial preoptic area (MPOA) of the mouse hypothalamus in response to lactation and hunger. Furthermore, we showed that heightened appetite in lactating mice was accompanied by increased activity of hunger-promoting agouti-related peptide (AgRP) neurons in the ARC. To assess the strength of hunger versus maternal drives, we designed a conflict assay where female mice chose between a food source or a chamber containing pups and nesting material. Although food-deprived lactating mothers prioritized parenting over feeding, hunger reduced the duration and disrupted the sequences of parenting behaviors in both lactating and virgin females. We discovered that ARC^AgRP^ neurons directly inhibit bombesin receptor subtype-3 (BRS3) neurons in the MPOA, a population that governs both parenting and satiety. Selective activation of this ARC^AgRP^ to MPOA^BRS3^ circuit shifted behaviors from parenting to food-seeking. Thus, hypothalamic networks are modulated by physiological states and work antagonistically during the prioritization of competing motivated behaviors.

---

In an ever-changing environment with limited resources, animals in the wild must learn to appropriately prioritize various needs to survive, reproduce, and propagate the species^10-13^. Different motivational systems in the brain are in place to navigate such needs. In situations where animals are faced with conflicting desires, motivational systems may function antagonistically in attempts of eclipsing one another, thereby promoting the behaviors that they govern^14-18^. Networks that are often distributed across the brain must therefore intersect to integrate multiple motivational signals, select which motivation to pursue, and ultimately engender a behavioral choice.

Two of the most fundamental drives are feeding and parenting — animals must procure food for sustenance and attend to the needs of newborn offspring. In female postpartum mammals, both feeding and parenting drives intensify concomitantly as lactation begins^1-5^. Producing milk puts animals in periods of negative energy balance, and as a result, lactating mothers exhibit increased food consumption^1,3,5^. Yet foraging for food cannot always take precedence, as abandoning helpless young in the nest is risky due to predation, starvation, and lack of warmth. This conflict is especially notable for most mammalian species, including the mouse *Mus musculus*, in which naturally occurring parental duties are shouldered solely by the mother^19,20^. Lactating mothers must therefore optimize when they seek food and provide maternal care, particularly in the wild when food sources are distant and sparse.

Past work has identified key neuronal populations underlying feeding and parenting behaviors, but these behaviors are often studied uncoupled from each other. Furthermore, while many studies have examined postpartum changes in circuits underlying parenting^3,4,21-23^, less is known about how hunger-promoting circuits adapt to accommodate the elevated energy requirements of milk production. Here, we describe how feeding and parenting neurons in the mouse hypothalamus are modulated by food-deprivation and lactation. By developing an ethologically relevant behavioral assay, we demonstrate how female mice resolve conflicts between feeding and parenting drives. We identify a novel population that regulates both parenting and satiety and delineate an intrahypothalamic circuit that underlies how hunger affects toggling between food-procurement and caring for pups. Our findings shed light on the neural coordination of essential survival behaviors and the balancing act carried out by animals in response to competing physiological and social incentives.

## Modulation of feeding behavior and circuits during lactation

To investigate how feeding behavior develops throughout different stages of motherhood, we measured the daily chow intake of singly housed female mice over several weeks. After a baseline measurement of virgin females, each female was co-housed with a male for two days. 14 out of 22 females became pregnant, and the remaining 8 females served as controls. While mice consumed 11.24 ± 0.27 kcal/day during baseline, consumption markedly surged to 41.99 ± 2.05 kcal/day shortly after parturition as mice began to lactate (Fig. 1a, Extended Data Fig. 1a). The mean daily food intake of lactating mice positively correlated with their litter size (Fig. 1b, Extended Data Fig. 1b-c), suggesting that the amount of milk produced to sustain pups necessitates a proportional increase in caloric demands. Indeed, halting lactation by weaning pups abruptly lowered food intake (Fig. 1a, Extended Data Fig. 1a-c).

**Fig. 1.**
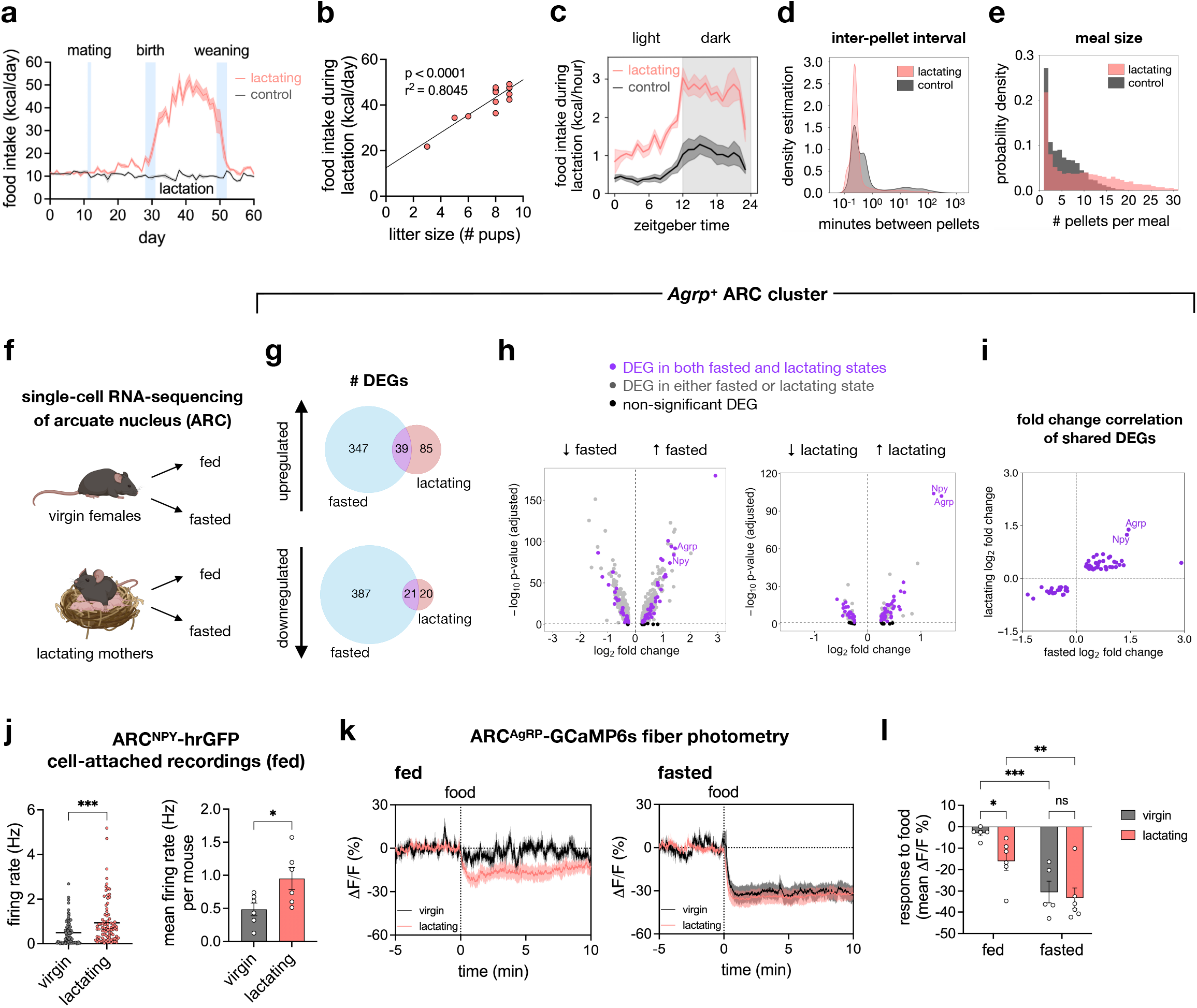
Lactating mice exhibit increased appetite and ARC^AgRP^ neuronal activity. **a**, food intake of lactating mice (*n =* 14) and female controls (*n* = 8). **b**, food intake during lactation (21 averaged days) vs. litter size (*n* = 14 lactating mice). **c-e**, FED3 measurements showing hourly food intake of 21 averaged days per mouse (**c**), inter-pellet intervals (**d**), and meal sizes (**e**) during lactation period (*n*_lactating_ = 5 mice, *n*_virgin_ = 6 mice). Pellets consumed within 60 s are part of one meal. **f**, schematic of 4 groups of mice for scRNA-seq created with BioRender. **g-i**, *Agrp*^+^ ARC cluster: Venn diagrams of DEGs in fasted and lactating states (**g**), volcano plots of DEGs comparing fasted virgins vs. fed virgins (left) and fed lactating mothers vs. fed virgins (right) (**h**), and fold change correlation of shared DEGs in fasted and lactating states (**i**). **j**, firing rate of ARC^NPY^-hrGFP neurons (left, *n*_virgin_ = 94 neurons, *n*_lactating_ = 88 neurons) and mean firing rate per mouse (right, *n* = 6 mice per group). **k**, fiber photometry recordings of ARC^AgRP^-GCaMP6s in fed and fasted virgin (*n* = 5) and lactating mice (*n* = 6). **l**, mean ΔF/F after food presentation. Plots show mean ± s.e.m. Data were analyzed using simple linear regression (**b**), unpaired two-tailed t-test (**j**), or 2-way repeated-measures analysis of variance (RM-ANOVA) with Fisher’s Least Significant Difference (LSD) test (**l**). \**P* < 0.05, \*\**P* < 0.01, \*\*\**P* < 0.001, \*\*\*\**P* < 0.0001. ns, non-significant. See Supplementary Table 1 for detailed statistical information.

While the increase in appetite during lactation has been previously described in rodents^5^, less is known about the circadian nature and microstructure of feeding during lactation. To gain higher temporal resolution of feeding behavior, we used Feeding Experimentation Devices version 3 (FED3)^24^ to dispense single 20-mg chow pellets, time-stamping each retrieval. Lactating mice displayed an increase in food intake in both dark and light cycles (Fig. 1c, Extended Data Fig. 1d-g). Furthermore, the inter-pellet intervals became shorter and meal sizes bigger during lactation, collectively resulting in higher food intake across circadian time (Fig. 1d-e).

To uncover the neural correlates of hyperphagia during lactation, we examined the arcuate nucleus of the hypothalamus (ARC), a critical brain region that regulates metabolism^6-8^. We performed single-cell RNA sequencing (scRNA-seq) of the ARC in 4 groups of female mice: virgins and lactating mothers that either had *ad libitum* food access (“fed”) or food-deprived overnight for 16-18 hours (“fasted”) (Fig. 1f). Analysis of 21,518 ARC neurons from the 4 groups revealed 34 genetically distinct clusters upon integration (Extended Data Fig. 2a-b). To identify which clusters were most responsive to fasting in terms of gene expression changes, we compared the fed and fasted virgin groups using principal component analysis (PCA) and calculated the contribution of each cluster to the first 8 principal components (PCs), which explained roughly 85% of the variance between the two groups. The cluster that contributed the most to the variance expressed *Agrp* (Extended Data Fig. 2c-e), encoding agouti-related peptide, which promotes hunger and feeding behavior^25,26^.

We thus focused on this *Agrp*^*+*^ cluster and analyzed the differentially expressed genes (DEGs) during fasting (comparing fasted virgins to fed virgins) and lactation (comparing fed lactating mothers to fed virgins). We identified hundreds of DEGs, 60 of which were differentially regulated in the same direction in both the fasted and lactating states (Fig. 1g-h, Supplementary Table 2). *Agrp* and *Neuropeptide Y* (*Npy)*, which encodes another potent orexigenic peptide that is co-released by ARC^AgRP^ neurons, showed similar degrees of upregulation during fasting and lactation (Fig. 1i). These results suggest that ARC^AgRP^ neurons exhibit a “transcriptomic hunger state” during lactation, even in fed animals.

The upregulation of *Agrp* and *Npy* and subsequent hyperphagia led us to hypothesize that ARC^AgRP^ neurons were more active during lactation. To test this hypothesis, we used an *Npy*-hrGFP line to label ARC^AgRP^ neurons and performed *ex vivo* cell-attached recordings in slice. Electrophysiological recordings from ARC^NPY^-hrGFP neurons showed an increase in firing rate in fed lactating mothers compared to fed virgins (Fig. 1j).

To corroborate that ARC^AgRP^ neurons are more active during lactation *in vivo*, we expressed a calcium indicator GCaMP6s in these cells and implanted a lens above the ARC for fiber photometry. Past studies have shown that ARC^AgRP^ are more active in fasted mice^27,28^, and bulk calcium imaging revealed that ARC^AgRP^ neurons are rapidly and sustainably inhibited by chow detection, specifically in fasted mice^29,30^. This suppression is not present in fed mice^29,30^, as ARC^AgRP^ neuronal activity is low in the fed state^27,28^. However, if ARC^AgRP^ neurons in fed lactating mothers are more active, then inhibition of ARC^AgRP^ calcium signals by chow should be observable even in this state. Indeed, presenting fed lactating mothers with chow significantly reduced ARC^AgRP^ calcium signals, an effect not seen in fed virgin females (Fig. 1k-l). Thus, both our *in vivo* and *ex vivo* studies demonstrate that ARC^AgRP^ neurons are more active during lactation.

### Transcriptomic modulation of MPOA by lactation and hunger

In addition to hyperphagia, the onset of motherhood in rodents is accompanied by an increased drive to provide maternal care^2-4^. The medial preoptic area (MPOA) is a crucial node in the hypothalamus that governs parenting behaviors^23,31-36^. Past work has shown that certain MPOA neurons undergo plasticity postpartum to accommodate the maternal needs of female mice^23^. To extend these past findings, we explored how MPOA neurons are transcriptionally modulated during lactation by performing scRNA-seq of the MPOA in the same groups of animals used for ARC scRNA-seq (Fig. 1f).

Analysis of 20,544 MPOA neurons revealed 40 genetically distinct clusters upon integration (Extended Data Fig. 3a-b). To determine which clusters are most sensitive to changes in their transcriptomic profiles during lactation, we compared the fed lactating and fed virgin groups using PCA. Cluster 3 contributed the most to the first 9 PCs when comparing the two groups (Fig. 2a), as can also be seen by its distance from the origin in PCA space (Fig. 2b). Interestingly, cluster 3 was the second largest contributor to the variance when comparing the fasted virgin and fed virgin groups (Fig. 2d-e). No other cluster displayed substantial changes in both the lactation and fasted states.

**Fig. 2.**
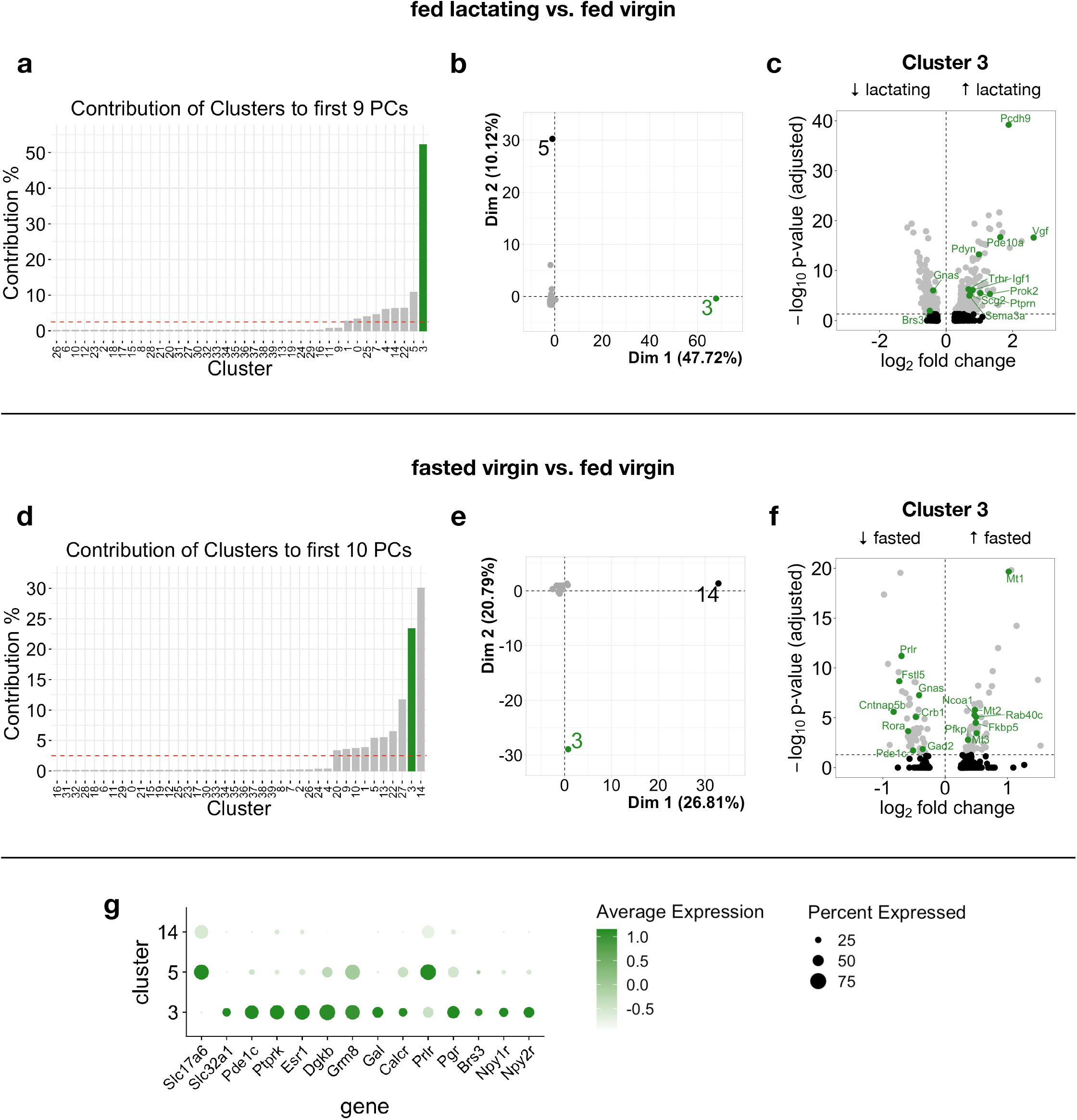
An MPOA neuronal cluster (#3) is highly transcriptionally responsive to both lactation and fasted states. **a-c**, comparison of fed lactating vs. fed virgin groups showing the contribution of each cluster to the first 9 PCs (**a**), magnitude of changes in transcriptomic profile of each cluster during lactation based on cluster’s distance from origin in PCA space (**b**), and DEGs in cluster 3 in the lactating state. **d-f**, comparison of fasted virgin vs. fed virgin groups showing the contribution of each cluster to the first 10 PCs (**d**), magnitude of changes in transcriptomic profile of each cluster during fasting based on cluster’s distance from origin in PCA space (**e**), and DEGs in cluster 3 in the fasted state (**f**). **g**, DotPlot of clusters 3, 5, and 14 showing expression of *Slc17a6 (Vglut2), Slc32a1 (Vgat)*, top 5 enriched genes in cluster 3 (*Pde1c, Ptprk, Esr1, Dgkb, Grm8*), parenting markers (*Gal, Calcr, Prlr, Pgr, Brs3)*, and NPY receptors (*Npy1r, Npy2r*). Dashed red line in **a** and **d** show contribution if all cluster contributed equally. Colors in **c** and **f** indicate select DEGs that have been implicated in parenting, appetite, endocrinology, metabolism, and other neuronal processes (green); other significant DEGs (gray); non-significant DEGs (black).

Many of the DEGs identified in cluster 3 during lactation or fasting have been implicated in parenting, appetite, endocrinology, metabolism, and other neuronal processes (Fig. 2c, f). The 5 most enriched genes in cluster 3 were *Pde1c, Ptprk, Esr1, Dgkb*, and *Grm8* (Fig. 2g). *Esr1* encodes estrogen receptor-alpha, and MPOA^Esr1^ neurons are important for parenting^34^. Other known parenting markers that were enriched in cluster 3 were *galanin (Gal)*^23,32,37^, *calcitonin receptor (Calcr)*^35^, *prolactin receptor (Prlr)*^33^, and *progesterone receptor (Pgr)*^23^ (Fig. 2g). Cluster 3 also expressed genes encoding NPY receptors, *Npy1r* and *Npy2r* (Fig. 2g), suggesting that this cluster may be sensitive to hunger states partly via an orexigenic NPY signaling pathway. These results demonstrate that the MPOA, particularly neuronal cluster 3, integrates signals related to both maternal care and energy balance.

### Conflict between feeding and parenting drives

Our finding that parenting neurons in the MPOA undergo pronounced transcriptional changes upon fasting suggests that hunger may influence parenting behaviors. Additionally, lactating mothers must carefully balance their increased appetite with their obligation to care for pups. To investigate how animals in different hunger states alternate between feeding and parenting behaviors, we designed a conflict assay in which female mice were placed in an arena with three chambers: one containing water and a FED3 as a food source (“feeding chamber”); one containing a shelter, dispersed pups, and scattered nesting material (“parenting chamber”); and an empty chamber in the middle. Lactating mothers and alloparenting virgins were tested under three different conditions: *fed (with pups), fasted (with pups)*, and *fasted (no pups)* (Fig. 3a).

**Fig. 3.**
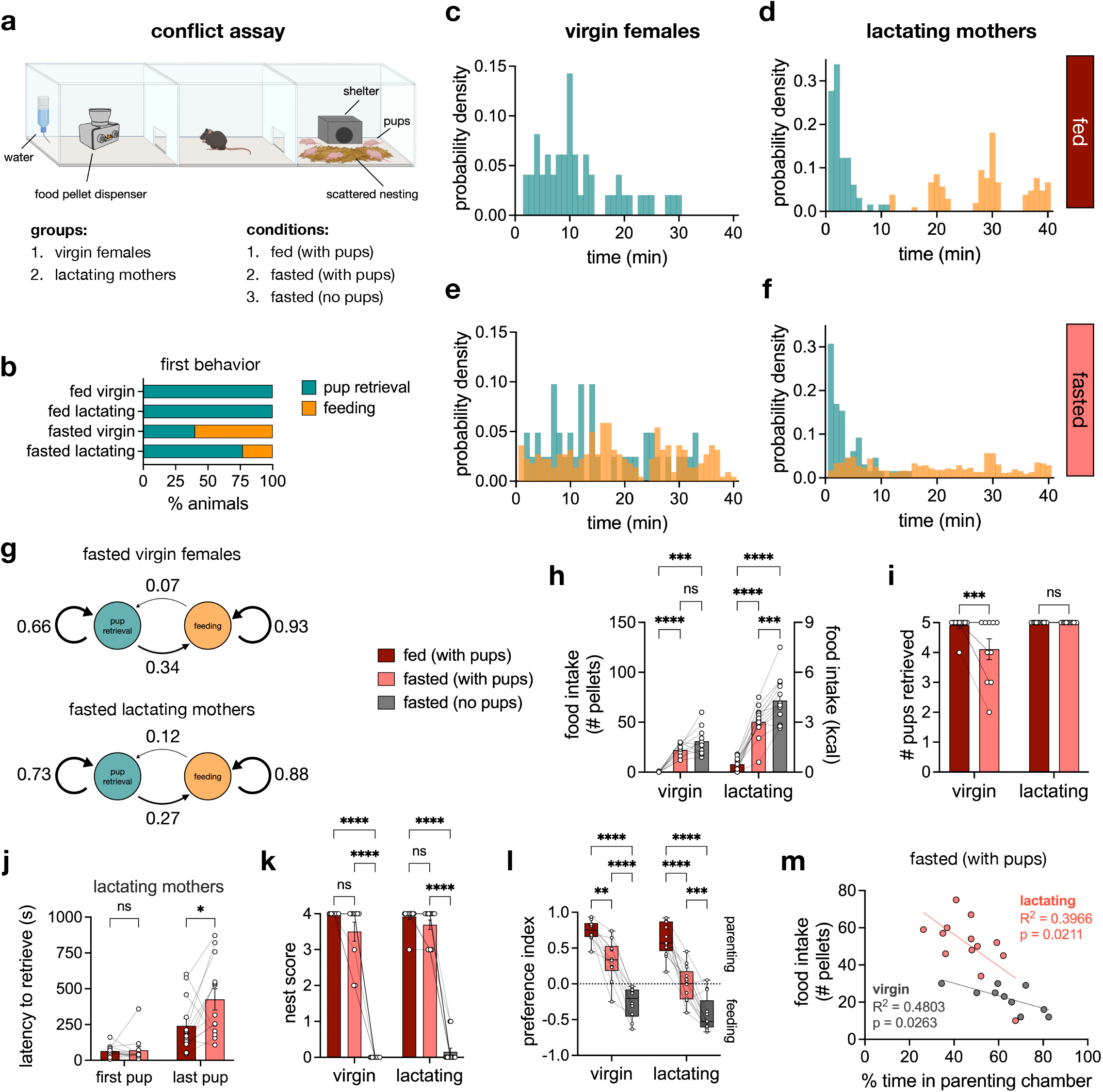
Hunger delays and reduces parenting behaviors in virgin females and lactating mothers. **a**, schematic of 40-minute conflict assay created with BioRender and adapted FED3 cartoon from a previous publication^19^. **b**, percent of animals that either performed pup retrieval or feeding as their first behavior (*n*_virgin_ = 10 mice, *n*_lactating_ = 13 mice). **c-f**, probability density (1-min bins) of pup retrieval and feeding events throughout the conflict assay (all trials). 1 total pellet consumed by 1 virgin female (fed, with pups) at time = 36.45 min is not shown to avoid skewing the visualization. **g**, transition probabilities between pup retrieval and feeding events in fasted virgin females and fasted lactating mothers. Only the feeding events prior to the completion of pup retrieval were used for the calculation. **h**, food intake. **i**, number of pups retrieved. **j**, latency to retrieve the first pup and last pup by lactating mothers. **k**, nest score. **l**, preference index based on time spent in parenting and feeding chambers. **m**, scatterplot of food intake (# pellets) vs. % time in parenting chamber of virgin and lactating females in the fasted (with pups) condition. Bar graphs show mean ± s.e.m. Box plots show median, upper/lower quartiles, and upper/lower extremes. Data were analyzed using two-way RM-ANOVA with Šidák’s multiple comparisons test (**h-l**) or simple linear regression (**m**). \**P* < 0.05, \*\**P* < 0.01, \*\*\**P* < 0.001, \*\*\*\**P* < 0.0001. ns, non-significant. See Supplementary Table 1 for detailed statistical information.

In the fed state, virgin and lactating mice began and completed pup retrieval to the shelter prior to feeding, establishing a priority for parenting (Fig. 3b-d, Extended Data Fig. 4a-b). However, in the *fasted (with pups)* condition when conflict was high, most virgins prioritized feeding while most lactating mothers prioritized pup retrieval (Fig. 3b, e-f, Extended Data Fig. 4c-d). The importance of parenting in lactating mothers could also be inferred from their significantly reduced food intake in the presence of pups even when fasted. In contrast, fasted virgins consumed comparable amounts of food regardless of pup presence (Fig. 3h). This effect was due to decreased maternal motivation in fasted virgins, as can be seen in the reduced number of pups retrieved (Fig. 3i). Although lactating mothers retrieved all pups, fasting delayed the completion of retrieval (Fig. 3j, Extended Data Fig. 4b, d). This deferral was a result of parenting interruption by sequences of feeding events in the fasted state, thus altering the transition probabilities between parenting and feeding behaviors (Fig. 3g, Extended Data Fig. 4b, d).

We also examined if hunger could affect maternal nest building. Fasting did not significantly affect the quality of the nest built in the presence of pups. However, virgin and lactating mice built little to no nest in the in the absence of pups, indicating that this action was geared toward pup care and not to regulate their own body temperature (Fig. 3k).

Using video-based animal tracking, the amount of time spent in each of the chambers was measured and used to calculate a preference index, where positive values indicated a preference for the parenting chamber, while negative values indicated a preference for the feeding chamber. Virgin and lactating mice preferred the parenting chamber in the *fed (with pups)* condition and the feeding chamber in the *fasted (no pups)* condition. However, the preference index moved closer to 0 in the *fasted (with pups)* condition, highlighting the conflict between parenting and feeding drives: pups reduced time spent in the feeding chamber, and reciprocally, hunger reduced time spent in the parenting chamber (Fig. 3l). Moreover, food intake in the *fasted (with pups)* condition was negatively correlated with the amount of time spent in the parenting chamber for both virgin and lactating mice, further exemplifying the mutual inhibition by parenting and feeding drives in a high-conflict context (Fig. 3m).

### Stimulating inhibitory ARC^AgRP^ projections to MPOA

The conflict assay demonstrated the capacity of hunger to reduce, delay, and disrupt the sequences of parenting behaviors. To gain entry to the neural circuitry underlying the selection between feeding and parenting behaviors, we manipulated ARC^AgRP^ neurons as they are GABAergic and inhibit satiety and other motivational systems^10,11,18,38-40^. Notably, ARC^AgRP^ neurons project to the MPOA^32-35,37,40^. Therefore, it is possible that the hunger-induced shift in priority from parenting to feeding in the conflict assay could be mediated by an inhibitory ARC^AgRP^ → MPOA circuit.

To test this possibility, we targeted channelrhodopsin-2 (ChR2) to ARC^AgRP^ neurons and implanted bilateral optic fibers above the MPOA to photostimulate ARC^AgRP^ → MPOA projections (Fig. 4a). Prior optogenetic studies have identified unique orexigenic ARC^AgRP^ projections^30,38,40,41^. To characterize the role of ARC^AgRP^ → MPOA in feeding, we measured food intake in fed virgin females without or with three different light stimulation paradigms: pre-stim (light on before food access), co-stim (light on during food access), or pre-& co-stim (light on before and during food access) (Fig. 4b). We found that even just pre-stimulation of ARC^AgRP^ → MPOA can elicit a significant increase in food intake, which was further heightened in the pre-& co-stim paradigm, independent of light intensity (Fig. 4c).

**Fig. 4.**
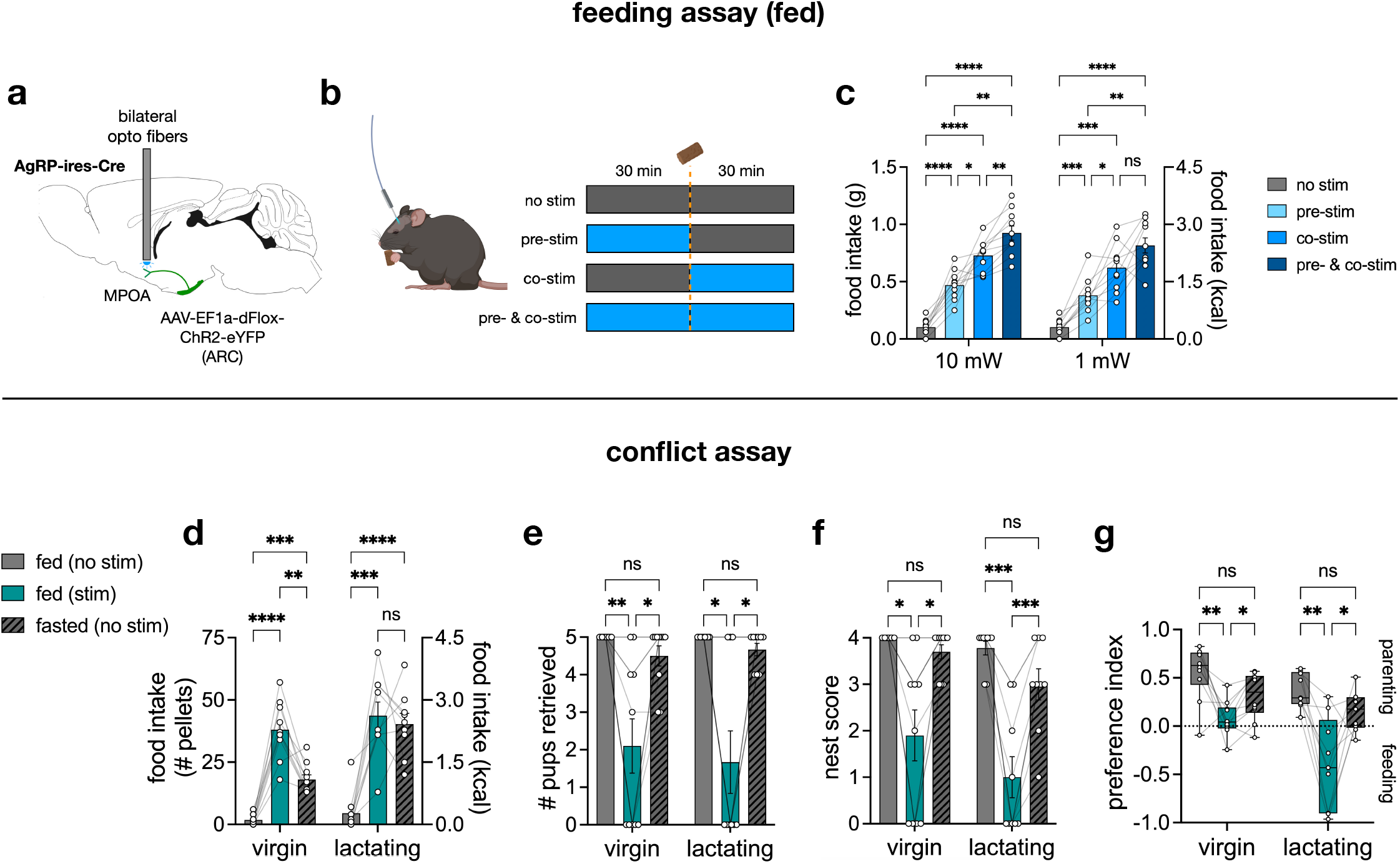
Optogenetic stimulation of ARC^AgRP^ projections to MPOA drives feeding and reduces parenting. **a**, sagittal view of the mouse brain (from Allen Brain Atlas) depicting the surgery procedure for optogenetics. **b**, schematic created with BioRender of stimulation paradigms in fed mice: no stim (light off), pre-stim (light on 30 min before food access), co-stim (light on 30 min during food access), and pre-& co-stim (light on 30 min before and 30 min during food access). **c**, 30 min food intake across different stimulation paradigms at 10 and 1 mW of light intensity; “no stim” datapoints are identical for 10 and 1 mW comparisons (*n* = 10 virgin females). **d-g**, results from the conflict assay showing food intake (**d**), number of pups retrieved (**e**), nest score (**f**), and preference index (**g**) (*n*_virgin_ = 10 mice, *n*_lactating_ = 9 mice; same subjects). Bar graphs show mean ± s.e.m. Box plots show median, upper/lower quartiles, and upper/lower extremes. Data were analyzed using two-way RM-ANOVA (**c**) or mixed-effects model analysis (**d-g**) with Šidák’s multiple comparisons test. \**P* < 0.05, \*\**P* < 0.01, \*\*\**P* < 0.001, \*\*\*\**P* < 0.0001. ns, non-significant. See Supplementary Table 1 for detailed statistical information.

Next, female mice were run in the conflict assay first as alloparenting virgins and later as lactating mothers with their own pups. The optogenetic conflict assay had three conditions with pups: *fed (no stim), fed (stim)*, and *fasted (no stim)*, representing no, artificial, and physiological hunger, respectively. ARC^AgRP^ → MPOA photostimulation increased feeding in virgins more than fasting did. However, the food intake of lactating mice was comparable in the *fed (stim)* and *fasted (no stim)* conditions (Fig. 4d). This result supports our earlier findings that ARC^AgRP^ neuronal activity is higher in lactating mothers than in virgins. Additionally, ARC^AgRP^ → MPOA photostimulation reduced pup retrieval, nest building, and veered preference toward the feeding chamber (Fig. 4e-g). Importantly, photostimulation of ARC^AgRP^-eYFP → MPOA had no effect on feeding or parenting (Extended Data Fig. 5). Thus, an ARC^AgRP^ → MPOA circuit can cause animals to prioritize food-seeking at the expense of parental duties.

### Inhibiting parenting neurons in MPOA

Since ARC^AgRP^ neurons are inhibitory, we hypothesized that they constrain the activity of parenting neurons in the MPOA during periods of hunger, swaying animals away from pup care in favor of food-seeking. We therefore aimed to silence parenting neurons in the MPOA to see if we could recapitulate a similar shift in behavior prioritization.

The MPOA is a molecularly and functionally heterogenous brain region, and as mentioned earlier, multiple populations have been shown to regulate parenting^32-35,37,42^. Targeting and inhibiting one of these labeled lines could have non-parenting effects, as MPOA^Esr1^ neurons, for example, are not only involved in parenting, but also other processes such as mating, metabolism, thermoregulation, and torpor^43-45^. Indeed, we found that chemogenetic inhibition of MPOA^Esr1^ reduced both parenting and feeding (Extended Data Fig. 6a).

To circumvent this heterogeneity, we employed the strategy Fos-Targeted Recombination in Activate Populations (FosTRAP)^46,47^ to label parenting-activated neurons in the MPOA with hM4Di, an inhibitory designer receptor exclusively activated by designer drugs (DREADD), agnostic of the neurons’ genetic markers (Fig. 5a). To this end, FosTRAP2 females were bilaterally injected with Cre-dependent hM4Di virus in the MPOA and mated post-surgery. TRAP was induced by injecting lactating mothers with 4-hydroxytamoxifen (4-OHT) after they had built a nest and retrieved their pups (“Parent-TRAP”). To control for cells activated by handling, a separate cohort of females underwent the same surgery but was injected with 4-OHT in the absence of pups or nest building (“Negative-TRAP”). Both Parent-TRAP and Negative-TRAP groups were then run in the conflict assay as lactating mothers with their own pups after injection of saline or clozapine N-oxide (CNO), a DREADD ligand.

**Fig. 5.**
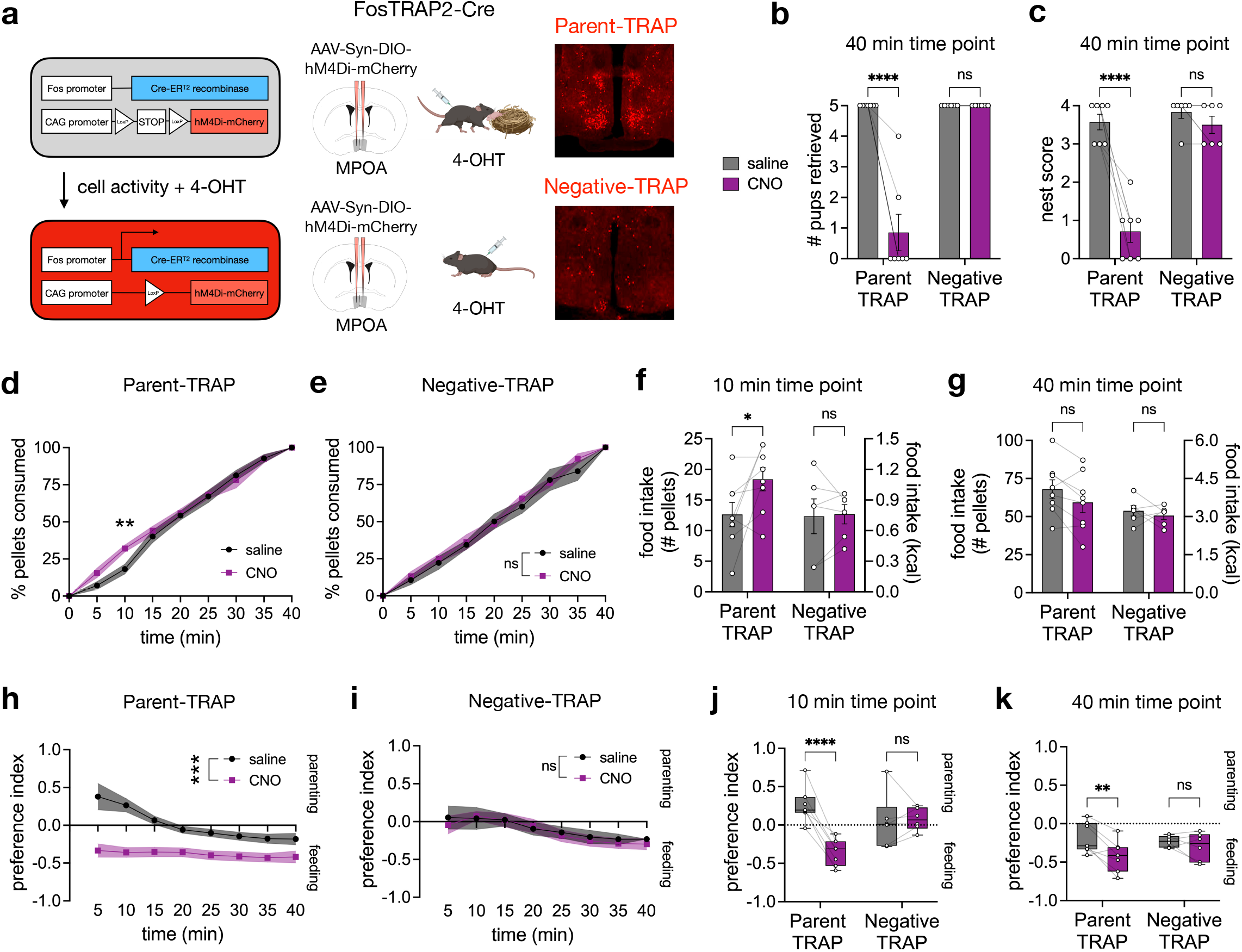
Inhibition of parenting neurons in MPOA reduces parenting and shifts priority to feeding in fasted lactating mothers. **a**, schematic of TRAP system and TRAP induction strategy to label parenting-responsive MPOA neurons with hM4Di-mCherry; created using BioRender and the Allen Brain Atlas. **b-k**, data from the conflict assay with fasted lactating Parent-TRAP (*n* = 7) and Negative-TRAP (*n* = 6) mice injected with saline or CNO. **b**, number of pups retrieved. **c**, nest score. **d-e**, percent of total pellets consumed over time in Parent-TRAP mice (**d**) and Negative-TRAP mice (**e**). **f-g**, food intake of mice at 10 minutes (**f**) and 40 minutes (**g**). **h-i**, preference index of Parent-TRAP mice (**h**) and Negative-TRAP mice (**i**) over time. Each datapoint represents the mean preference index from time = 0 min to that time point. **j-k**, preference index at 10 minutes (**j**) and 40 minutes (**k**). Bar and line graphs show mean ± s.e.m. Box plots show median, upper/lower quartiles, and upper/lower extremes. Data were analyzed using two-way RM-ANOVA with Šidák’s multiple comparisons test. Significance symbol in **d** is a post-hoc comparison of saline and CNO at time = 10 min. Significance symbols in **e, h, i** show the treatment effect of CNO. \**P* < 0.05, \*\**P* < 0.01, \*\*\**P* < 0.001, \*\*\*\**P* < 0.0001. ns, non-significant. See Supplementary Table 1 for detailed statistical information.

Inhibiting parenting neurons during the conflict assay significantly decreased pup retrieval and nest building in Parent-TRAP animals, an effect that was more pronounced in the fasted versus fed state, suggesting that hunger may provide further inhibition to parenting neurons, likely through the ARC^AgRP^ → MPOA circuit described above (Fig. 5b-c, Extended Data Fig. 7). In contrast, CNO injection had no effect on Negative-TRAP animals in either state (Fig. 5b-c, e-g, i-k, Extended Data Fig. 7). At an early time point (10 minutes), CNO injection caused fasted Parent-TRAP mothers to consume more food due to their minimal drive to prioritize maternal care (Fig. 5d, f). However, fasted Parent-TRAP mothers injected with saline eventually consumed comparable amounts of food as when they were injected with CNO (Fig. 5d, g), indicating that inhibition of MPOA parenting neurons merely shifted their priority to feeding rather than increased overall hunger levels. Indeed, saline-injected Parent-TRAP mothers initially preferred the parenting chamber but later switched preference for the feeding chamber (Fig. 5h). In contrast, CNO-injected Parent-TRAP mothers consistently preferred the feeding chamber when fasted, shirking their parenting responsibilities to eat (Fig. 5h, j-k). These results demonstrate that in a high-conflict context, MPOA parenting neurons in fasted lactating mothers regulate the prioritization of maternal care over food consumption, and silencing of these neurons can swing the priority.

### Interrogating the role of MPOA^BRS3^ neurons in parenting and satiety

The observation that inhibiting neurons in the MPOA shifts lactating mothers’ initial behaviors from parenting to feeding suggests the presence of a neuronal population that modulates both parenting and appetite. This population likely sits downstream of ARC^AgRP^ neurons. To pinpoint this putative cell-type, we looked at the genes expressed in *Fos*^+^ MPOA neurons in females after a parenting task. Multiplexed error-robust fluorescent *in situ* hybridization (MERFISH) of the MPOA^42^ identified two *Fos*^+^ clusters that had considerable expression of *Brs3*, which encodes bombesin receptor subtype-3. *Brs3* knock-out increases food intake and body weight^48^, as does delivering a BRS3 antagonist via intracerebroventricular injection, while BRS3 agonists reduce food intake and body weight^49^. Moreover, our sequencing data showed that 21% of neurons were *Brs3*^+^ in MPOA cluster 3, which was transcriptionally responsive to both lactation and fasting (Fig. 2g, Extended Data Fig. 3c-d). *Brs3* expression also colocalized with expression of known parenting markers (Extended Data Fig. 3e). Therefore, MPOA^BRS3^ neurons served as a candidate that could govern both parenting and satiety.

We expressed hM4Di in MPOA^BRS3^ neurons in alloparenting virgin females and ran the animals in our conflict assay to assess their role in behavior. Inhibiting MPOA^BRS3^ neurons considerably decreased pup retrieval and nest building both in the fed and fasted states (Fig. 6a-b). Notably, inhibiting MPOA^BRS3^ neurons reduced the preference index only in the fed but not the fasted state (Fig. 6c), suggesting that the shift in preference toward the feeding chamber may be in part due to increased sensation of hunger. Indeed, MPOA^BRS3^ neuronal inhibition increased food intake in the fed state in the conflict assay and in the animals’ home cages in the absence of pups (Fig. 6d-e). Thus, MPOA^BRS3^ neurons are important for regulating parenting and satiety.

**Fig. 6.**
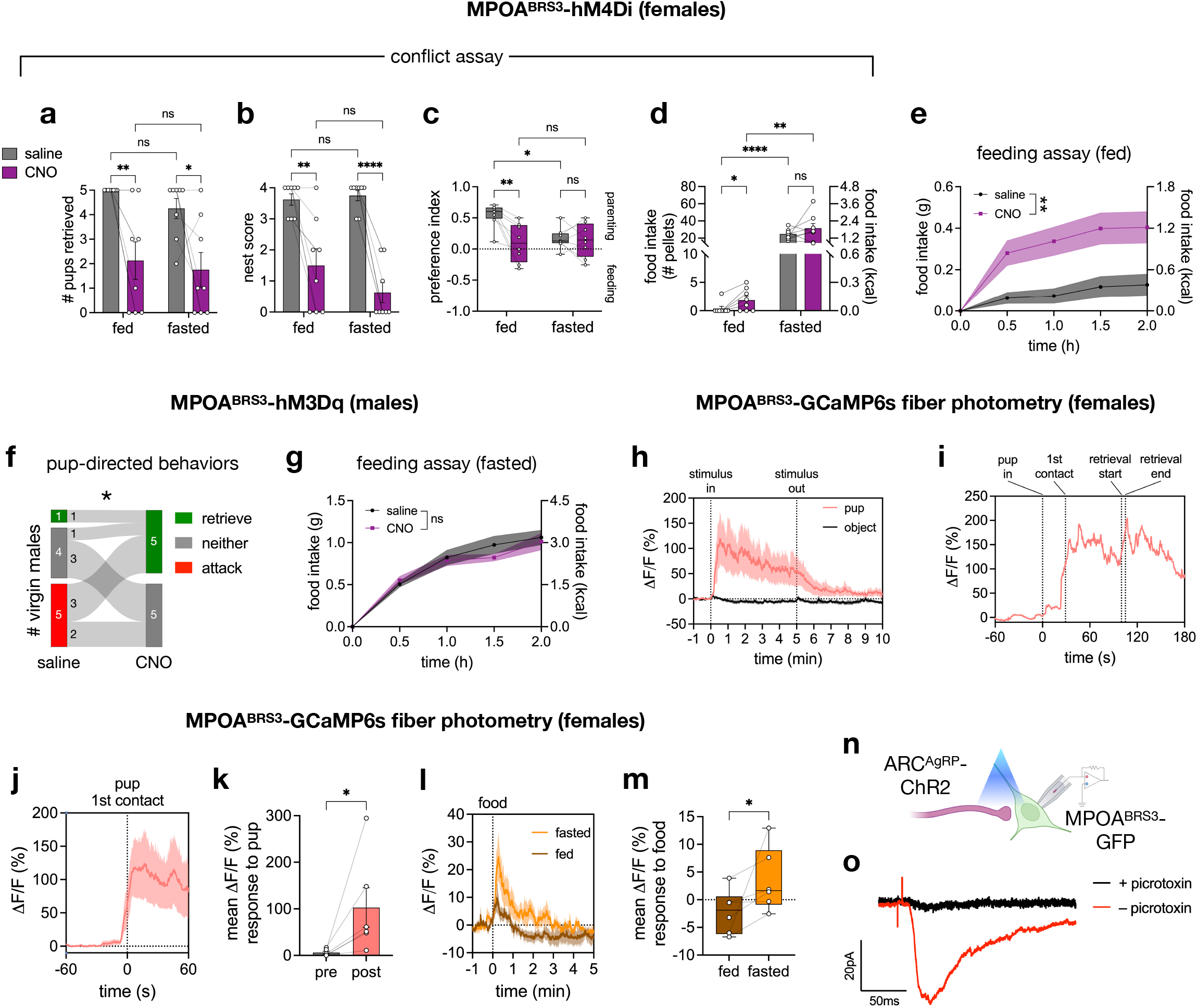
MPOA^BRS3^ neurons regulate parenting and feeding behaviors. **a-d**, MPOA^BRS3^-hM4Di virgin females (*n* = 8) run in the conflict assay showing number of pups retrieved (**a**), nest score (**b**), preference index (**c**), and food intake (**d**). **e**, food intake of fed MPOA^BRS3^-hM4Di virgin females (*n* = 7). **f**, number of MPOA^BRS3^-hM3Dq virgin males (*n* = 10) that retrieved or attacked pups, or neither. **g**, food intake of fasted MPOA^BRS3^-hM3Dq virgin males (*n* = 6). **h-m**, fiber photometry recordings of MPOA^BRS3^-GCaMP6s neurons in fed virgin females (*n* = 6) presented with either a pup or an object (**h**), representative female interacting with pup (**i**), signal aligned to 1st pup contact (**j**), mean signal pre- and post-pup contact (**k**), signal aligned to food access (**l**), and mean response to food in the fed and fasted states (**m**). **n**, circuit mapping strategy created with BioRender. **o**, averaged inhibitory post-synaptic current trace (32 trials) of a representative MPOA^BRS3^ neuron following light stimulation (red, no picrotoxin) and following light stimulation with picrotoxin (black). Bar and line graphs show mean ± s.e.m. Box plots show median, upper/lower quartiles, and upper/lower extremes. Data were analyzed using two-way RM-ANOVA (**a-e, g**) with Fisher’s LSD test (**a-d**), Chi-square test (**f**), or Wilcoxon matched-pairs signed rank test (**k, m**). Significance symbols in **e** and **g** show the treatment effect of CNO by two-way RM-ANOVA. \**P* < 0.05, \*\**P* < 0.01, \*\*\**P* < 0.001, \*\*\*\**P* < 0.0001. ns, non-significant. See Supplementary Table 1 for detailed statistical information.

Besides MPOA^Esr1^ and MPOA^BRS3^, we tested additional markers that are enriched in *Fos*^+^ MERFISH MPOA clusters^42^. Inhibiting MPOA^Calcr^ reduced pup retrieval only in the fasted state and had no effect on feeding. Inhibiting corticotropin-releasing hormone (MPOA^Crh^) or melanocortin-4 receptor (MPOA^MC4R^) neurons had no effect on feeding or parenting (Extended Data Fig. 6). Therefore, of all the populations we tested, MPOA^BRS3^ neurons may be the most relevant in underlying the toggling between parenting and feeding behaviors. We thus focused on further characterizing these neurons.

To see if activating MPOA^BRS3^ neurons can increase parenting drives, we expressed an excitatory DREADD hM3Dq in virgin males, which are typically non-parental. Virgin males expressing MPOA^BRS3^-hM3Dq were injected with either saline or CNO and then presented with pups in their home cages. Of the 10 saline-injected virgin males, 5 attacked pups, 1 retrieved pups, and 4 did neither. However, injecting the same animals with CNO completely abolished infanticide behavior and resulted in 5 virgin males retrieving pups, including most males that attacked when injected with saline (Fig. 6f). Therefore, activation of MPOA^BRS3^ neurons is sufficient to promote parenting in virgin males, and this effect is more pronounced than previous stimulations of other MPOA populations^32,35^.

To determine if activating MPOA^BRS3^ neurons can promote satiety and reduce feeding in the fasted state, MPOA^BRS3^-hM3Dq animals were fasted overnight, injected with either saline or CNO, and then given access to food. CNO injection had no effect on feeding (Fig. 6g). Thus, while MPOA^BRS3^ neuronal inhibition can reduce satiety and increase food intake in the fed state (Fig. 6d-e), activation is not sufficient to quell hunger in the fasted state, suggesting that the activity of hunger-promoting neurons may occlude chemogenetic activation of MPOA^BRS3^ neurons.

While *Fos* studies have implicated MPOA^BRS3^ neurons’ involvement in parenting^35,42^, it is not known when these neurons “come online” during different phases of parenting behaviors. To elucidate the temporal dynamics, we expressed GCaMP6s in MPOA^BRS3^ neurons in alloparenting virgin females to record calcium signals using fiber photometry. MPOA^BRS3^ neuronal calcium activity increased primarily as mice first made contact with the pup. The activity then remained elevated during and after retrieval as mice crouched over the pup in the nest (Fig. 6h-k). Single-cell microendoscopic calcium imaging of MPOA^BRS3^ neurons revealed a heterogenous response to pups, although most neurons were activated (Extended Data Fig. 8).

MPOA^BRS3^-GCaMP6s fiber photometry also showed a transitory increase in signal at the onset of food presentation, which was elevated in the fasted state (Fig. 6l-m). Recall that ARC^AgRP^ neurons are rapidly inhibited by the detection of food in fasted mice (Fig. 1k-l), and so the enhanced response of MPOA^BRS3^ neurons to food in the fasted state may be due to disinhibition by ARC^AgRP^ neurons. To test if ARC^AgRP^ neurons directly inhibit MPOA^BRS3^ neurons, ChR2 was expressed in ARC^AgRP^ neurons while GFP was expressed in MPOA^BRS3^ neurons (Fig. 6n). Brains were sliced for *ex vivo* whole-cell recordings of MPOA^BRS3^-GFP neurons with tetrodotoxin in the bath to block action potentials. Stimulating ARC^AgRP^-ChR2 axons with blue light evoked time-locked inhibitory post-synaptic currents in MPOA^BRS3^-GFP neurons in 4 female mice, which was abolished by the addition of picrotoxin, a GABA_A_ receptor antagonist (Fig. 6o). Thus, ARC^AgRP^ neurons monosynaptically inhibit MPOA^BRS3^ neurons via GABA signaling. This ARC^AgRP^ → MPOA^BRS3^ circuit may therefore underlie the process by which hunger can redirect an animal’s motivation from caring for pups to foraging for food.

## DISCUSSION

Our study delineates how physiological states such as hunger and lactation modulate the gene expression and activity of ARC and MPOA neurons, offering insights into their roles in regulating behavioral priorities. We confirm previous findings that ARC^AgRP^ neurons adapt to lactation’s increased energy demands by upregulating *Agrp* and *Npy*^50-52^. Using scRNA-seq, we extend these observations by identifying the differential regulation of 58 additional genes associated with both fasting and lactation, opening new avenues of research into their roles in lactation-induced hyperphagia. In addition to gene expression changes, we show that ARC^AgRP^ neurons undergo plasticity during lactation by increasing their firing rate and magnifying calcium responses to food. Further work is required to investigate how various factors, such as pup sensory cues and the changing hormonal landscape, can remodel ARC^AgRP^ neuronal properties.

To understand how female mice balance feeding and maternal drives, we designed a behavioral paradigm that pits these drives against each other. We found that although lactating mothers have the propensity to prioritize maternal care even when fasted, hunger can reduce and disrupt overall parenting behaviors in both virgin and lactating females. Photostimulation of ARC^AgRP^ → MPOA mimics these hunger-induced effects, while acute silencing of Parent-TRAP neurons in the MPOA shifts fasted mothers’ priority toward feeding. Using circuit mapping, we uncover that ARC^AgRP^ neurons monosynaptically inhibit MPOA^BRS3^ neurons, a population that we discovered has an important dual role in parenting and appetite, despite constituting only 6% of all MPOA neurons. Our findings support a neural circuit mechanism whereby hunger signals activate ARC^AgRP^ neurons which in turn inhibit MPOA^BRS3^ neurons, thus promoting food-seeking and abrogating parental drives.

It has been previously posited that shared circuitry between parenting and satiety systems may exist to aid the survival of offspring by reducing the drive of mothers to forage for food, thereby encouraging them stay with their offspring^53^. We provide evidence supporting this notion. First, our MPOA sequencing dataset points to a neuronal cluster that exhibits profound transcriptional changes during motherhood and periods of fasting. This cluster expresses several parenting markers, suggesting an integrative function of the MPOA in regulating maternal care and energy balance. Future work can probe how specific genes within this cluster may contribute to this function. Second, and more specifically, we demonstrate that inhibition of MPOA^BRS3^ neurons reduces parenting and satiety, particularly in the fed state, causing animals to favor feeding. We also show that MPOA^BRS3^ neuronal activity rapidly increases during parenting and upon food presentation. The response to food is augmented in the fasted state, possibly encoding an anticipatory satiety signal.

One of the well-studied roles of MPOA^BRS3^ neurons is their ability to raise body temperature upon activation by cold exposure^54,55^. In rats and mice, pups increase ultrasonic vocalizations in cold environments, such as outside of the nest^56,57^. It is possible that MPOA^BRS3^ neuronal activation signals coldness in both oneself and others, such as pups. This function is reminiscent of “mirror neurons” in the ventromedial hypothalamus that are activated both by observing and executing social aggression^58^. The repurposing of MPOA^BRS3^ neurons may be a strategy of the nervous system to efficiently drive cold-defense and parenting behavior. Thus, MPOA^BRS3^ neurons may serve as a junction in a multi-pronged system that regulates the physiological needs of both parents and offspring.

Altogether, our work provides a comprehensive characterization of how ARC and MPOA neurons function in hunger and lactation states. We developed an ethologically relevant behavioral assay to investigate the interplay between feeding and parenting drives, demonstrating how mice manage their physiological need for nutrients against the social incentive to nurture pups. We identified an intrahypothalamic neural circuit that orchestrates the prioritization of these fundamental behaviors, thus deepening our understanding of how such drives are regulated in response to changing physiological states. Our findings offer insights that could inform therapeutic strategies for psychiatric disorders, such as postpartum depression, potentially aiding individuals in developing a stronger bond with their infants and better prioritizing their needs.

## METHODS

### Animals

Mouse husbandry and experimental procedures were approved by the National Institutes of Health Animal Care and Use Committee. Mice were singly housed (unless used for breeding) in 22-24°C with a 12-hour light/12-hour dark cycle. Mice were fed with *ad libitum* standard chow (Teklad F6 Rodent Diet 8664) and water, unless food-deprived overnight for 16-18 hours for fasted experiments. Mice were 8-20 weeks old during the time of surgeries and experiments. The following mouse lines were used (with strain number from Jackson Laboratory): C57BL/6 (000664), *Agrp-*ires*-*Cre (012899), *Npy-*hrGFP (006417), *Fos-*2a*-*iCre^ERT2^ “TRAP2” (030323), *Esr1-*2a*-*Cre (017911), *Brs3-* ires*-*Cre (030540), *Calcr-*ires*-*Cre (037028), *Crh-*ires*-*Cre (012704), *Mc4r-*2a*-*Cre (030759). To generate the *Brs3-*T2A*-*FlpO line, the *Brs3-FlpO* allele was constructed on a C57BL/6 background with a Gly-Ser-Gly (GGAAGCGGA) linker, 54-bp T2A sequence^59^, and 1299-bp FlpO sequence^60^ inserted into the *Brs3* stop codon using CRISPR technology (Cyagen Biomodels LLC, Santa Clara, CA). Two synonymous mutations (Ala392, GCC to GCA; Glu396, GAG to GAA) were also introduced for CRISPR. Mice were genotyped by PCR (annealing 60°C, extension 72°C) using primers x475 (WT forward): 5’-GGGAGTGCACATGTCTCTGA-3’, x564 (common reverse): 5’-CATGAACAACAGAAAACCCAGA-3’, and x803 (FlpO forward): 5’-CCCATCAGCAAGGAGATGAT-3’, which generate 171 bp (WT) and 273 bp (FlpO) products.

### Food intake measurements

For daily chow intake measurements, chow was weighed every day for singly housed female mice, except for the 2 days with co-habituation with a male. For FED3 measurements, female mice were housed in PhenoTyper cages (Noldus). 2 FED3s (Open Ephys) with synchronized clocks set in “free feeding mode” were placed in each cage and checked daily for jamming. FED3s were filled with 20-mg food pellets (5-TUM, TestDiet) and replenished daily to maximum capacity. Food pellet retrievals in the light and dark cycles were determined based on the timestamps. Inter-pellet intervals and meal sizes were calculated using the software FED3 Viz and the associated code in Python. Pellets were part of the same meal if they were retrieved within 60 s of each other.

### Single-cell RNA-sequencing sample collection

4 groups (fed virgins, fasted virgins, fed lactating mice, fasted lactating mice) of 14-week-old female C57BL/6 mice (*n* = 3 mice pooled per group) were used for scRNA-seq. The litter size for the lactating groups were 8 pups and pups were 13-15 days old at the time of sample collection. Mice were anesthetized with isoflurane and then rapidly sacrificed by decapitation. Brains were extracted and sliced at 225-μm thickness using a vibratome as previously described^61^. Briefly, slices were taken in an ice-cold high-sucrose slicing solution consisting of the following components (in mM): 87 NaCl, 75 sucrose, 25 glucose, 25 NaHCO_3_, 1.25 NaH_2_PO_4_, 2.5 KCl, 7.5 MgCl_2_, 0.5 CaCl_2_ and 5 ascorbic acid saturated with 95% O_2_/5% CO_2_. Slices were then enzyme-treated for ∼15 min at 34°C with protease XXIII (2.5 mg/mL; Sigma) in a high-sucrose dissociation solution containing the following (in mM): 185 sucrose, 10 glucose, 30 Na_2_SO_4_, 2 K_2_SO_4_, 10 HEPES buffer, 0.5 CaCl_2_, 6 MgCl_2_, 5 ascorbic acid (pH 7.4) and 319 mOsm. Slices were washed three times with cold dissociation solution then transferred to a trypsin inhibitor/bovine serum albumin (TI/BSA) solution (10 mg/mL; Sigma) in cold dissociation solution. 7-8 slices containing the MPOA and 9-10 slices containing the ARC were obtained per group. MPOA and ARC were microdissected using ultra fine forceps and prepared for sequencing as previously described^62^. Briefly, microdissected tissue was kept in cold TI/BSA dissociation solution until trituration. Immediately before dissociation, tissue was incubated for ∼10 min at 37°C, then triturated with a series of small bore fire-polished Pasteur pipettes. Cell-suspension was kept on ice until single-cell capture. Cell viability was measured for each sample preparation using a Countess II automated cell counter (ThermoFisher) and was between the range of 67.5% to 82.3% before processing with the 10x Genomics platform^63^. 12,000 cells were loaded for capture onto an individual lane of a Chromium Controller (10x Genomics) for each sample. Single-cell capture, barcoding, and library preparation were performed using the 10x Chromium platform according to the manufacturer’s protocol (#CG00052) using version 3 (V3) chemistry.

### Single-cell RNA-sequencing bioinformatics analysis

ARC and MPOA data were analyzed and clustered separately using the R Package Seurat (https://satijalab.org/seurat/)^64^. Cells from the 4 groups of female mice were integrated using Reciprocal PCA and normalized using SCTransform. Cells that were classified as neurons using well-established neuronal marker genes via DropViz (http://dropviz.org)^65^ were used for clustering. Different resolutions for clustering were tested by calculating the silhouette score and examining the resulting UMAP, and 0.5 was chosen for both ARC and MPOA datasets. Clusters were then filtered based on quality control metrics (mitochondrial/erythrocyte gene expression and sequencing saturation), high expression of select neuronal markers, and low expression of select non-neuronal markers. DEGs and cluster marker genes were identified using the Wilcoxon Rank Sum test. Initial clustering at 0.5 resolution revealed two *Agrp*^*+*^ ARC clusters that displayed a high degree of similarity, and they were later combined into a single cluster for subsequent analyses. To determine the contribution of each cluster to the variance between two groups of mice (e.g., fed lactating mothers vs. fed virgins, or fasted virgins vs. fed virgins), DEGs for each cluster with an adjusted p-value < 0.05 and a log_2_ fold change > 1.5 were identified and compiled into a matrix of log_2_ fold changes and used as input for PCA. The contribution of each cluster to the first number of PCs explaining ∼85% of the variance was then calculated.

### Behavioral conflict assay

Pups used for all experiments were 5 days old or younger. Mice were singly housed in PhenoTyper cages with a dark, opaque plastic shelter (10 cm length x 10 cm width x 5.7 cm height) containing two holes (3.2 cm diameter) that allow for entry. Nesting material (crinkle paper) was placed inside the shelter. Mice were fed using a FED3 and regular chow in the bedding. To promote alloparenting in virgin females, mice were exposed to pups for 2-3 consecutive days. To generate lactating mothers, female mice were co-housed with males, which were removed until the females were visibly pregnant (∼7 days before parturition). Alloparenting virgin females and lactating mothers were habituated for 2 consecutive days in the conflict assay 3-chamber arena made of transparent plexiglass (72 cm length x 40 cm width x 22 cm height, with 2 partition walls dividing the length to create 3 equal-sized chambers; door openings in partition walls are 5 cm width x 9 cm height). For lactating mothers, the 1st day of habituation was the day they gave birth. The parenting chamber contained scattered nesting material, 5 dispersed pups, and a shelter containing 1 pup and 3-5 strands of nesting material. The feeding chamber contained a FED3 and 50 mL falcon tube with water and a lick spout. For habituation, mice were first placed in the empty middle chamber with the doors closed, which were then opened by the experimenter after ∼5 min and mice were allowed to explore the arena for 1 hour. Habituation was repeated the following day for 1 hour with the sides of the parenting and feeding chambers switched. Mice for chemogenetics experiments were also habituated to saline intraperitoneal (i.p.) injections. For experiments, the 3-chamber arena was set up and mice were placed in the empty middle chamber for 2 min before opening the doors. Video was recorded through EthoVision XT 16 (Noldus) at 30 frames per second using a webcam. Zones were created on EthoVision to track which chamber the mouse was occupying at all time points. For frames in which the mouse was undetected (i.e. inside the shelter), the location was replaced with the last detected location. The tracking data was analyzed by calculating the time spent in each zone/chamber. A preference index was calculated based on the time spent in the parenting and feeding chambers with the following formula: (parenting - feeding)/(parenting + feeding). Pups were considered retrieved once they were brought inside the shelter (scored manually). A qualitative nest score was given with the following rubric: 0 (no nesting material was brought into the shelter), 1-2 (some nesting material was brought into the shelter, but the nest was flat), 3 (over half of the nesting material was brought into the shelter and the nest was concave), 4 (all of the nesting material was brought into the shelter and the nest was concave). Experiments were completed within the first 6 hours of the light cycle. The order of the experimental conditions was randomized.

### Stereotaxic surgery and viral injections

Surgeries were performed as previously described^18^. Briefly, mice were anesthetized with 2-5% isoflurane first in a chamber then mounted on a stereotaxic apparatus. A heating pad set at 37°C was placed below the mice. Mice were subcutaneously injected with 1 mg/kg buprenorphine and 1 mL saline. The skull was leveled and craniotomy was performed. Viruses were injected at a speed of 25 nL/min using a glass pipette with a tip diameter of 20–40 μm attached to an air pressure system, and the pipette was withdrawn 10 min after injection. The following coordinates relative to Bregma were used: ARC (A/P -1.47, M/L ±0.25, D/V - 5.7), MPOA (A/P +0.1, M/L ±0.33, D/V -5.4). See sections below for viruses used and volume. Mice were allowed to recover for at least 3 weeks post-surgery before further experiments.

### Optogenetics

*Agrp-*ires*-*Cre virgin females were bilaterally injected with AAV9-EF1alpha-dFlox-hChR2-eYFP (Addgene 20298) or AAV1-EF1alpha-DIO-eYFP (Addgene 270560) (300 nL) in the ARC. Optogenetic cannulae (200 μm diameter core; BFH37-200 Multimode; NA 0.37; Thor Labs) were implanted bilaterally above the MPOA (A/P +0.1, M/L ±0.33, D/V -5.1), with one fiber tilted at 10 degrees to increase the distance between the two implants above the skull. Fibers were set onto the skull using C&B Metabond Quick Cement. Metabond was allowed to completely dry, then dental cement was applied for extra stability. For behavioral experiments, fiber optic patch cords (1 m long, 200 μm diameter; Doric Lenses) attached a laser via a commutator (Doric Lenses) were attached to fiber optic cannulae with zirconia sleeves (Doric Lenses). Open-loop pulse trains (20 Hz; 2 s ON, 2 s OFF; 50% duty cycle; 473 nm; Laserglow laser technologies) were programmed using a waveform generator (PCGU100; Valleman Instruments). Light intensity was measured and set at 1 or 10 mW. For the feeding assay, virgin females first habituated to the set-up for at least 3 consecutive days. Experiments with fed mice were begun during the first 4 hours of the light cycle. Mice received no photostimulation, or photostimulation before, during, or before and during food access (each mouse received all 4 conditions; the order was randomized). Food intake was measured 30 min after chow presentation. For the conflict assay, alloparenting virgin females and lactating mothers were first habituated to being hooked up to the patch cords in the three-chamber arena for 2 consecutive days. The laser was turned on at 10 mW with open-loop pulse trains (program described above) for 10 min before the start of the assay and remained on throughout the entire 40-min duration. The design of the conflict assay was as described above, except the partition walls were modified and part of the shelter’s top was removed to allow mice to freely move while attached to the patch cords.

### TRAP induction

4-hydroxytamoxifen (4-OHT, Sigma H6278) was dissolved in 55°C DMSO at a concentration of 4 mg/mL and stored in 25 μL aliquots in -20°C. On the day of experiments, aliquots were thawed and heated to 55°C. 500 μL saline solution with 2% Tween 80 was also heated to 55°C, which was then added to the 4-OHT aliquot and vortexed immediately. 475 μL saline was added to obtain a final volume of 1 mL working solution, which was kept at 55°C until right before injections. 10 μL working solution per 1 g of mouse was i.p. injected, resulting in a concentration of 10 mg/kg injected into the mouse. Injection was done during the first 4 hours of the light cycle. For Parent-TRAP, TRAP2 virgin females were injected bilaterally with AAV9-hSyn-DIO-hM4Di-mCherry (Addgene 44362) (200 nL) in the MPOA then mated with a male 1 week post-surgery. Males were removed when females were visibly pregnant. Females were then housed in PhenoTyper cages with their nest inside the shelter. Females were habituated to handling and i.p. injection for 4 consecutive days starting on the day they give birth. On the 5th day, pups and nesting material was scattered in the cages (with 1 pup remaining inside the shelter) and mice were allowed to retrieve pups to the shelter and build a nest. 1 hour later, mice were i.p. injected with 4-OHT, and the pups and nesting material were re-scattered to allow mice to display maternal care again and re-activate parenting neurons. 1 week later, the Parent-TRAP animals were co-housed with a male and were tested in the conflict assay with their 2nd litter. For Negative-TRAP, surgery was performed in TRAP2 virgin females as described above and were i.p. injected with 4-OHT 3 weeks post-surgery without any pup stimulus or nest building assay. 1 week later, Negative-TRAP animals were co-housed with males and tested in the conflict assay after giving birth.

### Chemogenetics

CNO (Tocris 4936) was dissolved in saline at 6 mg/mL, aliquoted, and kept frozen in -20°C. On the day of experiments, aliquots were thawed and diluted in saline to 0.3 mg/mL. Mice were i.p. injected with 3 mg/kg CNO and experiments were run 1 hour afterward. For the conflict assay, the 1st experimental day for the TRAP lactating mothers was always a fasted condition, followed by two days of fed conditions, and ending with a fasted condition. This was to ensure that all habituation and experiments would be completed within 5 days after the pups were born. The TRAP lactating mothers either received saline or CNO injection on the first experimental day, and the injection was alternated for the next 3 experimental days. For alloparenting virgin females, the order of the conditions in the conflict assay was more randomized. Virgin females (*Esr1*-ires-Cre, *Brs3*-ires-Cre, *Calcr*-ires-Cre, *Crh*-ires-Cre, *Mc4r*-ires-Cre) for the conflict assay were injected bilaterally with AAV9-hSyn-DIO-hM4Di-mCherry (200 nL) in the MPOA. Virgin males (*Brs3-* ires*-*Cre) were injected bilaterally with AAV9-hSyn-DIO-hM3Dq-mCherry (Addgene 44361) (50 nL) in the MPOA. For parenting assays, virgin males were injected with saline or CNO. 1 hour after injection, 3 pups were placed in their home cage outside of the nest for 30 min. Males were classified by their pup-directed behaviors: retrieved at least 1 pup to the nest, attacked at least 1 pup (i.e. pup has at least 1 bite mark), or neither. For feeding assays, female and male mice were injected with saline or CNO within the first 4 hours of the light cycle and placed in new cages with new nesting material. 1 hour after injection, chow was added to the home cage and weighed every 30 min for 2 hours to measure food intake.

### Fiber photometry

AAV1-Syn-Flex-gCaMP6s-WPRE (Addgene 100845) (250 nL) was injected unilaterally in the ARC for *Agrp-* ires*-*Cre females or MPOA for *Brs3-*ires*-*Cre females. A 400-μm optic-fiber cannula was implanted above the ARC (A/P -1.47, M/L +/-0.25, D/V -5.6) or MPOA (A/P +0.1, M/L +/-0.33, D/V -5.1) and secured with MetaBond and dental cement. Photometry recordings were performed with Tucker-Davies Technologies system as previously described^18^. Data used for analysis were collected at 8 Hz. Video was also recorded with frames aligned to specific timestamps of photometry data collection. The mice used were alloparenting virgin females or lactating mothers. Mice were hooked up to the patch cord, placed in a new cage with nesting material from their home cage, and habituated for at least 20 min prior to the start of recording. For *Agrp-*ires*-*Cre recordings, a pup was placed with each fed mouse. A baseline measurement was taken, and recording continued after the addition of chow. For *Brs3-*ires*-*Cre recordings, a baseline measurement was taken, and recording continued after the addition of a pup, an object, or chow. Percent changes in calcium fluorescence signal were calculated using the following formula: ΔF/F = (F_raw_ - mean)/mean x 100, where “F_raw_” is the raw fluorescence value and “mean” is the mean fluorescence value 5 minutes before chow presentation (for *Agrp-*ires*-*Cre photometry recordings) or 1 minute before stimulus presentation (for *Brs3-* ires*-*Cre photometry recordings).

### Microendoscope imaging

AAV1-Syn-Flex-gCaMP6s-WPRE diluted 1:4 in saline was injected (250 nL) unilaterally in MPOA. Then, a 25-guage hollow guide needle was implanted slowly (∼1 mm/min) above MPOA (A/P +0.1, M/L +/- 0.33, D/V -5.1). After removing the guide needle, a ProView Integrated GRIN lens (0.5 mm diameter, 8.4 mm length; Inscopix) was implanted slowly (∼1 mm/5 min) at A/P +0.1, M/L +/-0.33, D/V -5.3. Metabond was applied to fill the space between the skull and the baseplate. Dental cement was applied to cover Metabond. Mice were allowed to recover for at least 4 weeks post-surgery before screening for calcium signal. Inscopix nVista miniscope and Inscopix nVision behavioral camera were synchronized for calcium imaging and behavioral video acquisition. Videos were collected at 20 Hz. LED power and gain of the miniscope were adjusted so that the right tail of the signal histogram was at ∼40%. Lens focus was adjusted to find the plane with the most in-focus cells. After mice had been habituated to the Inscopix set-up for at least 3 days, experiments were run in which a baseline measurement was taken followed by the addition of a pup, an object, or chow. Data were analyzed using Inscopix Data Processing Software. Miniscope videos pre-processed, downsampled spatially by 4 and temporally by 2, and motion-corrected. Neurons were identified by taking the maximum intensity projection of the entire video and manually drawing regions of interest (ROI) to calculate fluorescence values in ROI’s over time. Traces for each neuron were calculated using the formula as described above for fiber photometry. The baseline period was 1 minute before stimulus presentation.

### Electrophysiology

Cell-attached recordings were made in the ARC of female *Npy-*hrGFP reporter mice (virgin or lactating 14-16 days postpartum; age-matched littermates). Brain slices were obtained and stored at 30°C in a heated, oxygenated chamber containing (in mM) 124 NaCl, 4.4 KCl, 2 CaCl_2_, 1.2 MgSO_4_, 1 NaH_2_PO_4_, 10.0 glucose, and 26.0 sodium bicarbonate in artificial cerebrospinal fluid (aCSF) before being transferred to a submerged recording chamber maintained at 30°C (Warner Instruments). Recording electrodes (2.8-3.8 MΩ) were pulled with a Flaming-Brown Micropipette Puller (Sutter Instruments) using thin-walled borosilicate glass capillaries. Cell-attached spiking was measured in current-clamp mode using electrodes filled with aCSF, while maintaining a loose membrane seal. Average spike frequency was analyzed in the last 2 min of a 5-min recording. For channelrhodopsin-assisted circuit mapping, *Brs3-*T2A*-*FlpO; *Agrp-*ires*-*Cre female mice received bilateral injections of AAV8-EF1a-fDIO-GCaMP6s (Addgene 105714) (250 nL) in the MPOA and AAV9-EF1alpha-dFlox-hChR2-mCherry (Addgene 20297) (300 nL) in the ARC. Brain slices were acquired (see above) and inhibitory postsynaptic currents were measured upon light stimulation (2.5 mW blue LED power, 5 ms pulse) in voltage-clamp mode using electrodes (3-5 MΩ) filled with an intracellular recording solution containing (in mM) 130 CsCL, 10 HEPES, 10 EGTA, 2 ATP, 0,2 GTP, pH to 7.4, 290-295 mOsmol. 3 mM Kynurenic acid was included in the aCSF bath and neurons were held at −70 mV to isolate light-evoked GABAergic synaptic transmission. 500 nM tetrodotoxin and 100 μM 4-aminopyridine were also included in the aCSF bath. During the last recording of each slice, inhibitory GABA currents were confirmed by washing on 25 µmol/L picrotoxin. Signals were acquired via a Multiclamp 700B amplifier (Molecular Devices), digitized at 20kHz, filtered at 3kHz, and analyzed using Clampfit 11.4 software (Molecular Devices).

### Statistics

Error bars, exact *n* values, and statistical tests are described in the figure legends. Statistical tests were run using Python, R, or Prism. 2-4 cohorts of combined sample sizes of 5-14 mice were used for all electrophysiological and behavioral experiments. Each replicate was one mouse (or one neuron for electrophysiological recordings) that was run once in the experiment. Data normality was assumed except for in Fig. 6k, 6m, and Extended Data Fig. 5e-f. Exact *p*-values, *t* values, *F* values, and degrees of freedom are provided in Supplementary Table 1.

## OTHER INFORMATION

### Data availability

Data are provided in all main and extended figures. Source data, including a link to the scRNA-seq dataset, will be uploaded on: https://github.com/ivancalcantara/FeedingParenting/

### Code availability

Code used to analyze scRNA-seq data was derived from: https://satijalab.org/seurat/

Code used to analyze FED3 data was derived from: https://github.com/earnestt1234/FED3_Viz

Custom codes for our study will be uploaded on https://github.com/ivancalcantara/FeedingParenting/

## Acknowledgments

We thank Alexander Fleischmann, Wei Li, W. Scott Young, and members of the Michael J. Krashes, Marc L. Reitman, and Andrew Lutas labs for discussions and support; and Glenn Parham for assisting with coding and data analysis. This research was supported by the Intramural Research Program of the National Institutes of Health, the National Institute of Diabetes and Digestive and Kidney Diseases (DK075088, DK075087-06), and the NextGen Sequencing Grant from the National Institute of Environmental Health Sciences.

## Author contributions

I.C.A. and M.J.K. conceived the project and designed the experiments. I.C.A., C.L., L.E.M., C.M.M., I.d.A.S., E.O.K., A.I.G., R.A.P., and S.R.G. performed the experiments and analyzed the data. I.C.A., C.G., B.N.P., and J.L.L. analyzed the scRNA-seq data. C.X. generated the *Brs3-* T2A*-*FlpO mouse line. I.C.A. and B.N.P. made the figures. I.C.A. wrote the manuscript, which all authors reviewed and edited.

## Ethics declaration

The authors declare no competing interests.

---

**Extended Data Fig. 1.**
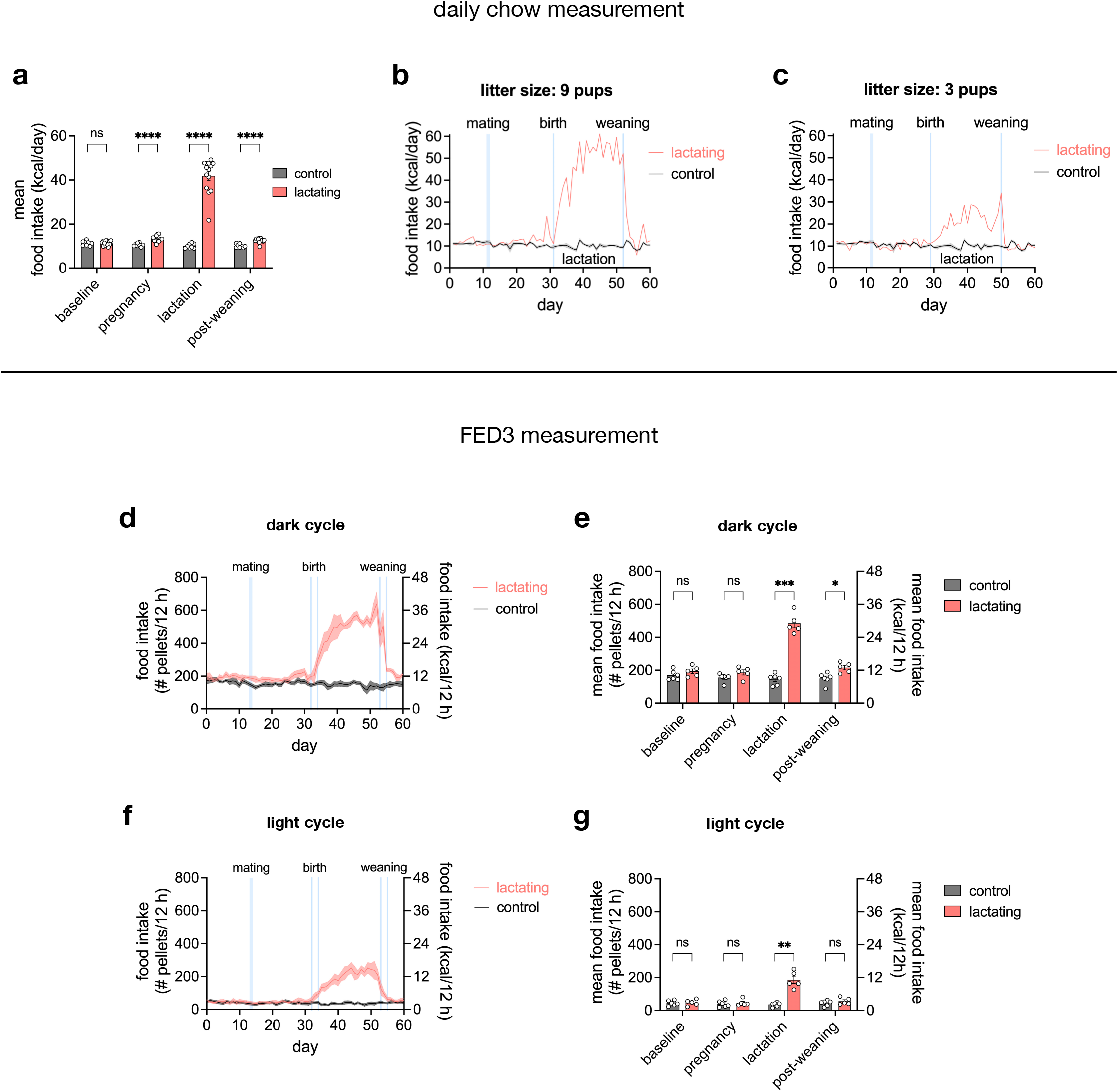
Lactating mice exhibit increased appetite during dark and light cycles. **a**, mean daily food intake of control (*n* = 8) and lactating (*n* = 14) mice during baseline, pregnancy, lactation, and post-weaning. **b-c**, daily food intake of representative lactating mice with 9 pups (**b**) or 3 pups (**c**). **d-g**, daily food intake measurement using FED3 in the dark cycle (**d-e**) and light cycle (**f-g**) of control (*n* = 6) and lactating (*n* = 5) mice during baseline, pregnancy, lactation, and post-weaning. Plots show mean ± s.e.m. Data were analyzed using two-way RM-ANOVA with Šidák’s multiple comparisons test (**a, e, g**). See Supplementary Table 1 for detailed statistical information.

**Extended Data Fig. 2.**
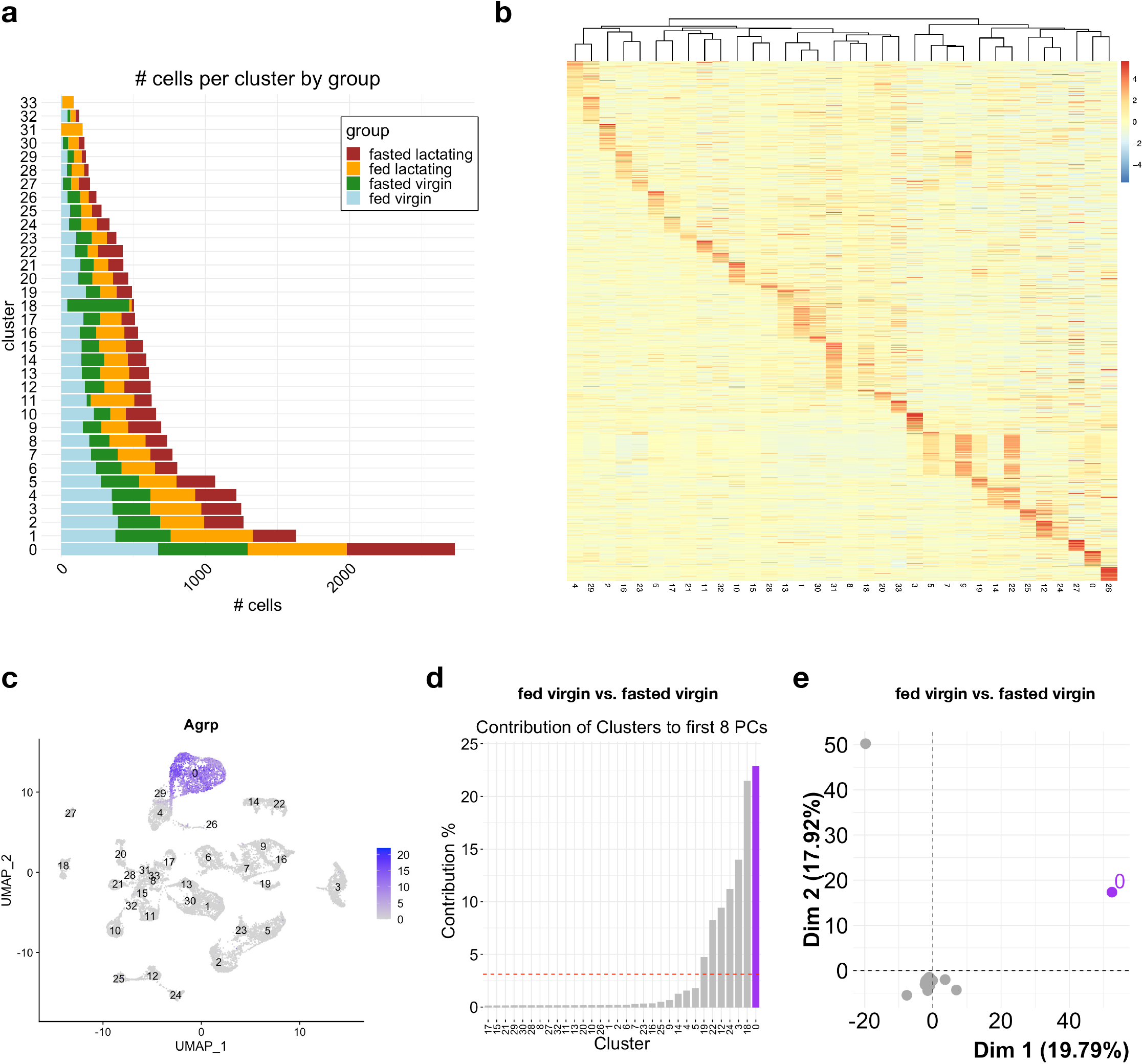
*Agrp*^*+*^ neuronal cluster in the ARC exhibits the most changes in gene expression upon fasting. **a**, number of cells per ARC neuronal cluster split by group. **b**, heat map of relative expression of marker genes in each ARC neuronal cluster with dendrogram (top) showing similarities between clusters. **c**, FeaturePlot showing ARC neurons and clusters in UMAP space with *Agrp* expression localized in cluster 0. **d**, contribution of ARC clusters to first 8 PCs of the variance in gene expression between fasted virgin and fed virgin groups, highlighting *Agrp*^*+*^ cluster 0. Dashed red line shows contribution if all cluster contributed equally. **e**, visualization of how much ARC clusters change transcriptionally between fasted virgin and fed virgin groups based on clusters’ distance from the origin in PCA space, highlighting *Agrp*^*+*^ cluster 0.

**Extended Data Fig. 3.**
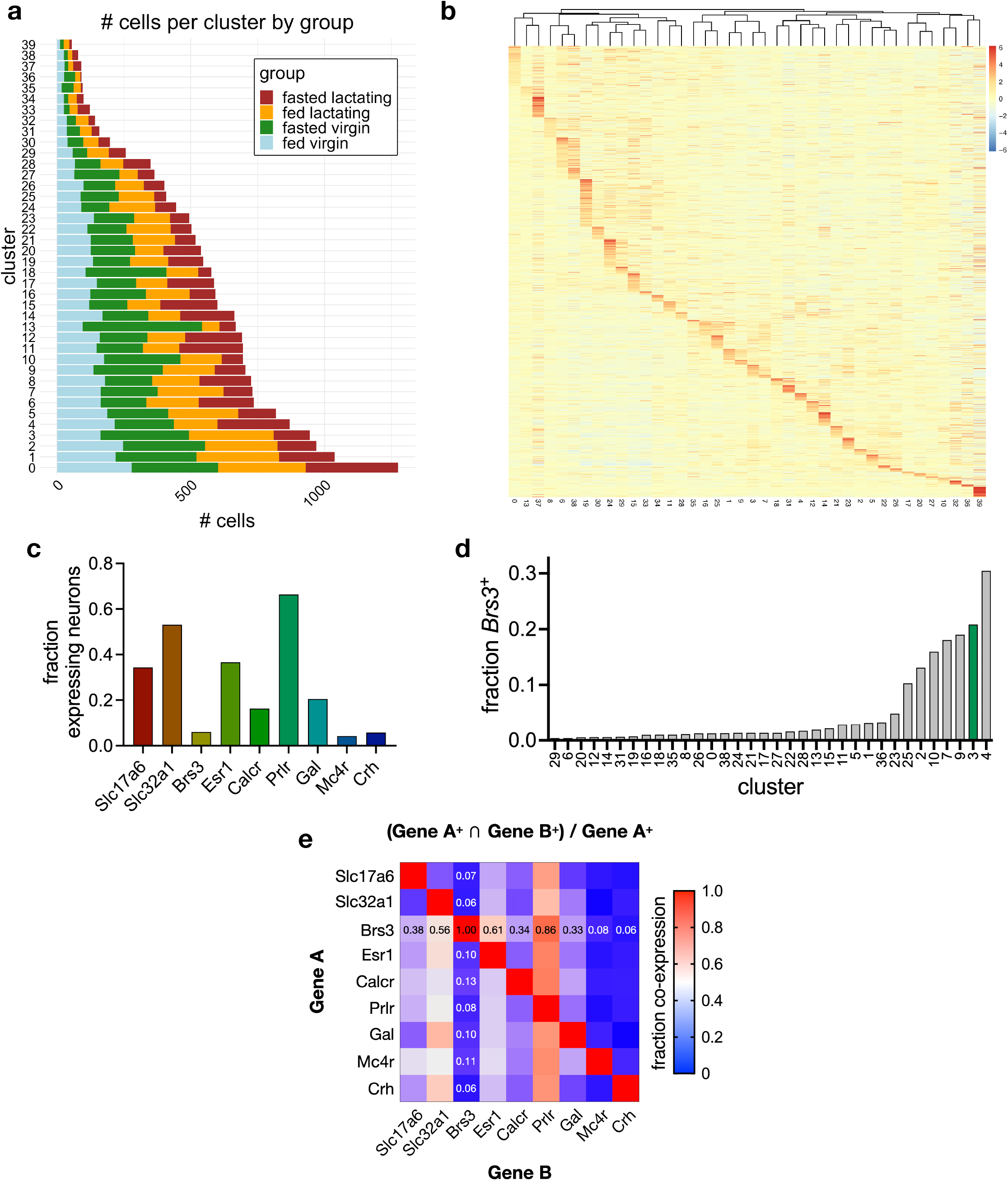
MPOA neuronal clusters co-express several genetic markers for parenting-promoting neurons. **a**, number cells per MPOA neuronal cluster separated by group. **b**, heat map of relative expression of marker genes in each MPOA neuronal cluster with dendrogram (top) showing similarities between clusters. **c**, fractions of MPOA neurons that are positive, i.e. contains at least 1 unique molecular identifier (UMI), for select genes. **d**, fractions of neurons that are *Brs3*^*+*^ per MPOA cluster, highlighting cluster 3 in green. **e**, co-expression matrix showing fraction of Gene A^+^ MPOA neurons that are also Gene B^+^, highlighting co-expression with *Brs3* (e.g., 38% of *Brs3*^*+*^ neurons are *Slc17a6*^*+*^).

**Extended Data Fig. 4.**
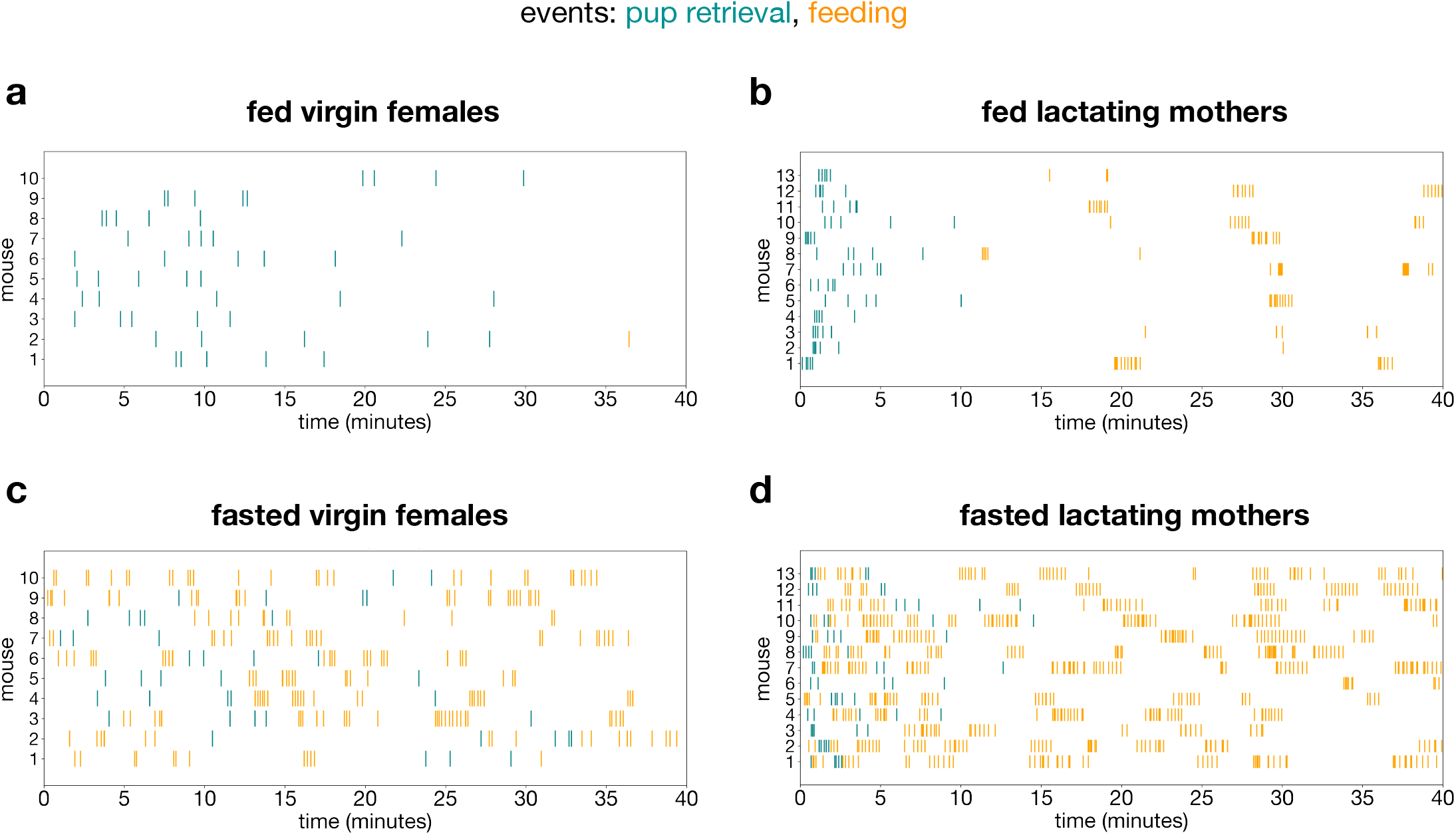
Lactation and fasted states affect pup retrieval and feeding behavioral sequences. **a-d**, raster plots showing pup retrieval (blue) and feeding (orange) events in fed virgin females (**a**), fed lactating mothers (**b**), fasted virgin females (**c**), and fasted lactating mothers (**d**) (*n*_virgin_ = 10 mice, *n*_lactating_ = 13 mice).

**Extended Data Fig. 5.**
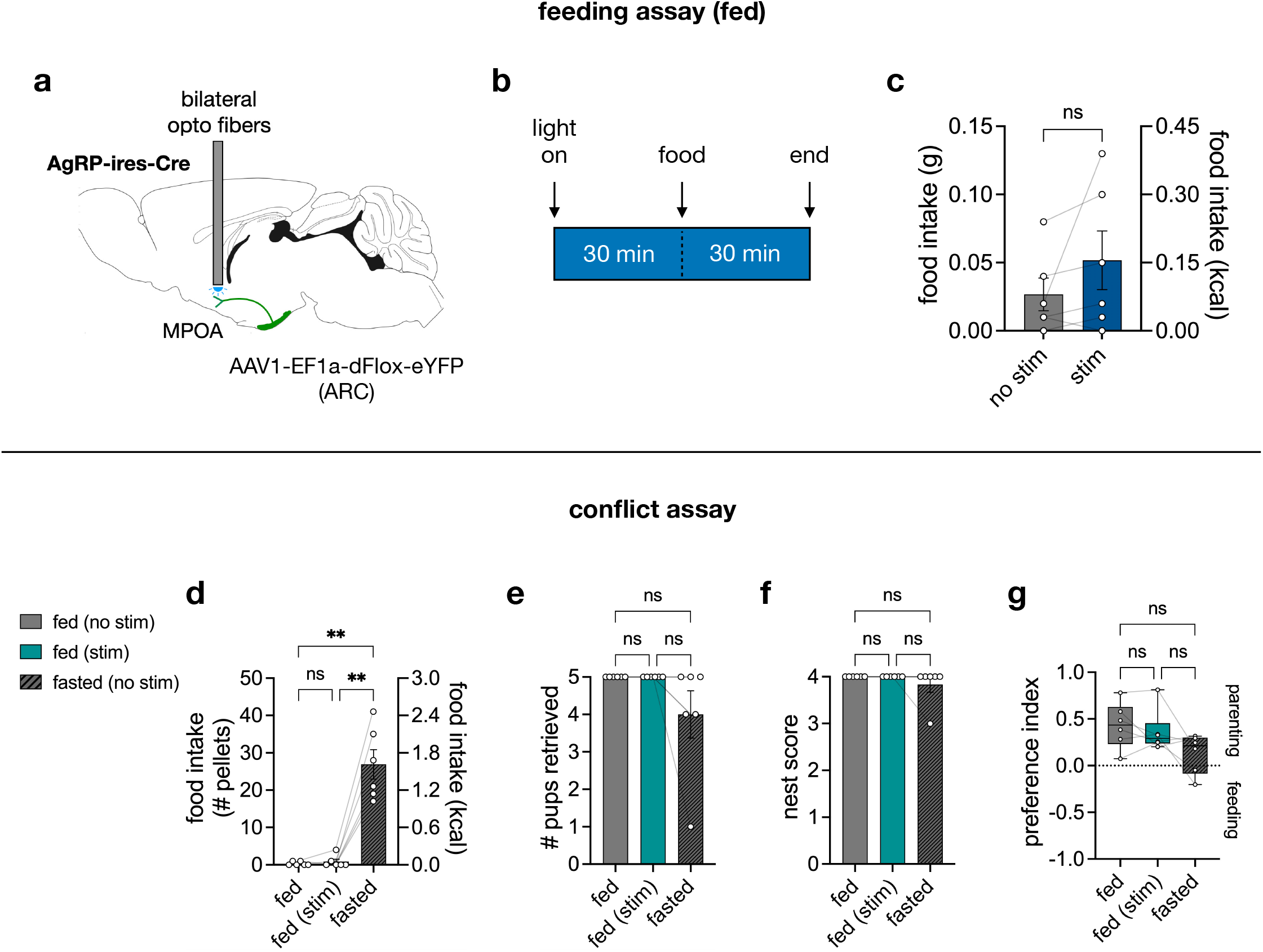
Photostimulation of ARC^AgRP^-eYFP projections to MPOA does not affect feeding or parenting behaviors. **a**, sagittal view of the mouse brain (Allen Brain Atlas) depicting the surgery procedure for optogenetics. **b**, schematic of photostimulation paradigm in fed mice for feeding assay (*n =* 6 virgin females). **c**, 30 min food intake with or without photostimulation. **d-g**, data from conflict assay showing food intake (**d**), number of pups retrieved (**e**), nest score (**f**), and preference index (**g**) (*n =* 6 virgin females). Bar graphs show mean ± s.e.m. Box plots show median, upper/lower quartiles, and upper/lower extremes. Data were analyzed using a two-tailed paired t-test (**c**), one-way RM-ANOVA with Tukey’s multiple comparison’s test (**d, g**), or Friedman test with Dunn’s multiple comparisons test (**e-f**). \**P* < 0.05, \*\**P* < 0.01, \*\*\**P* < 0.001, \*\*\*\**P* < 0.0001. ns, non-significant. See Supplementary Table 1 for detailed statistical information.

**Extended Data Fig. 6.**
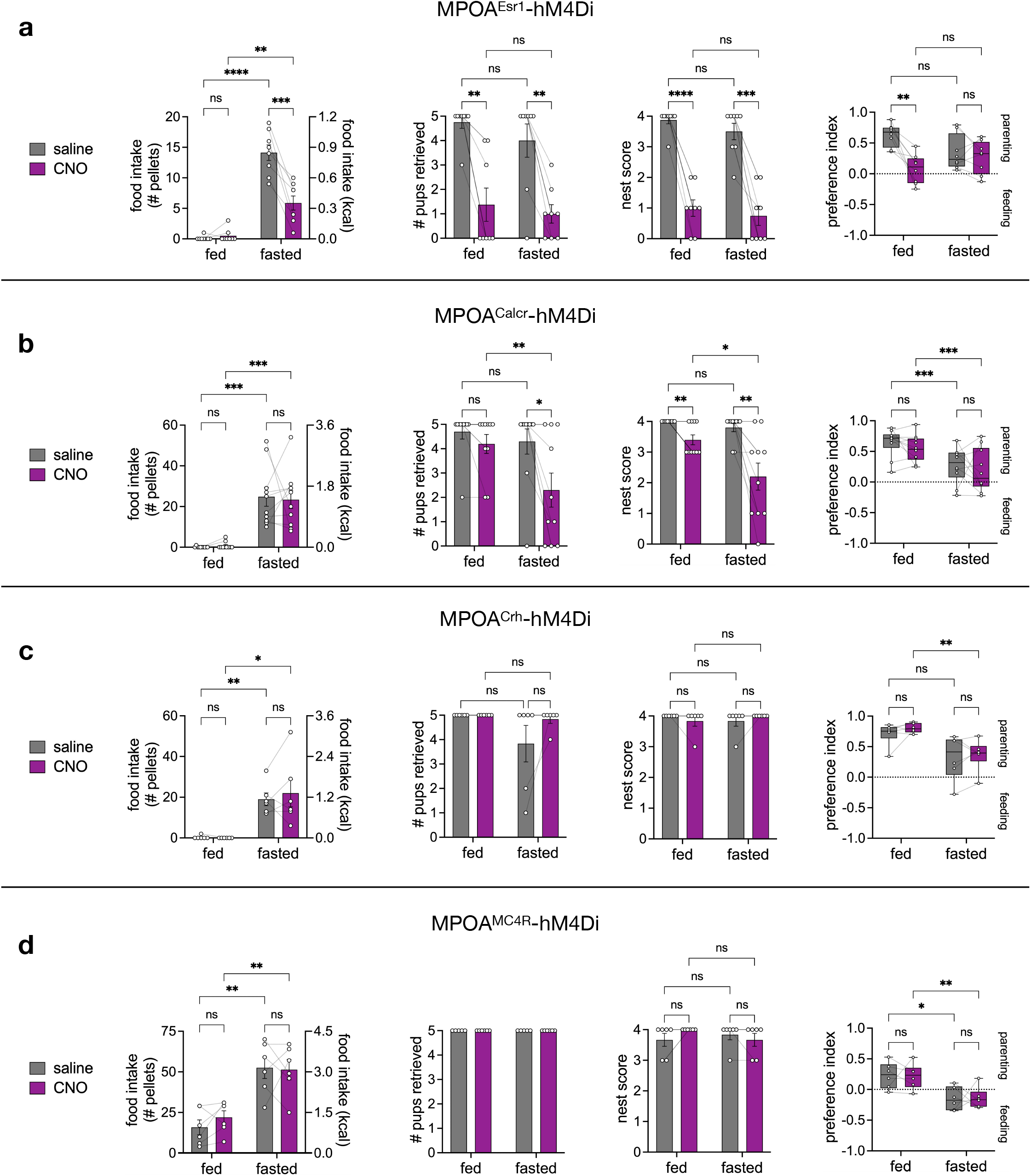
Inhibition of Esr1, Calcr, Crh, or MC4R neurons in MPOA with hM4Di. **a-d**, data from conflict assay showing (from left to right) food intake, number of pups retrieve, nest score, and preference index of animals expressing hM4Di in MPOA^Esr1^ (**a**, *n* = 8 virgin females), MPOA^Calcr^ (**b**, *n* = 10 virgin females), MPOA^Crh^ (**c**, *n* = 6 virgin females), or MPOA^MC4R^ (**d**, *n* = 6 lactating mothers). Bar graphs show mean ± s.e.m. Box plots show median, upper/lower quartiles, and upper/lower extremes. Data were analyzed using two-way RM-ANOVA with Fisher’s LSD test. See Supplementary Table 1 for detailed statistical information.

**Extended Data Fig. 7.**
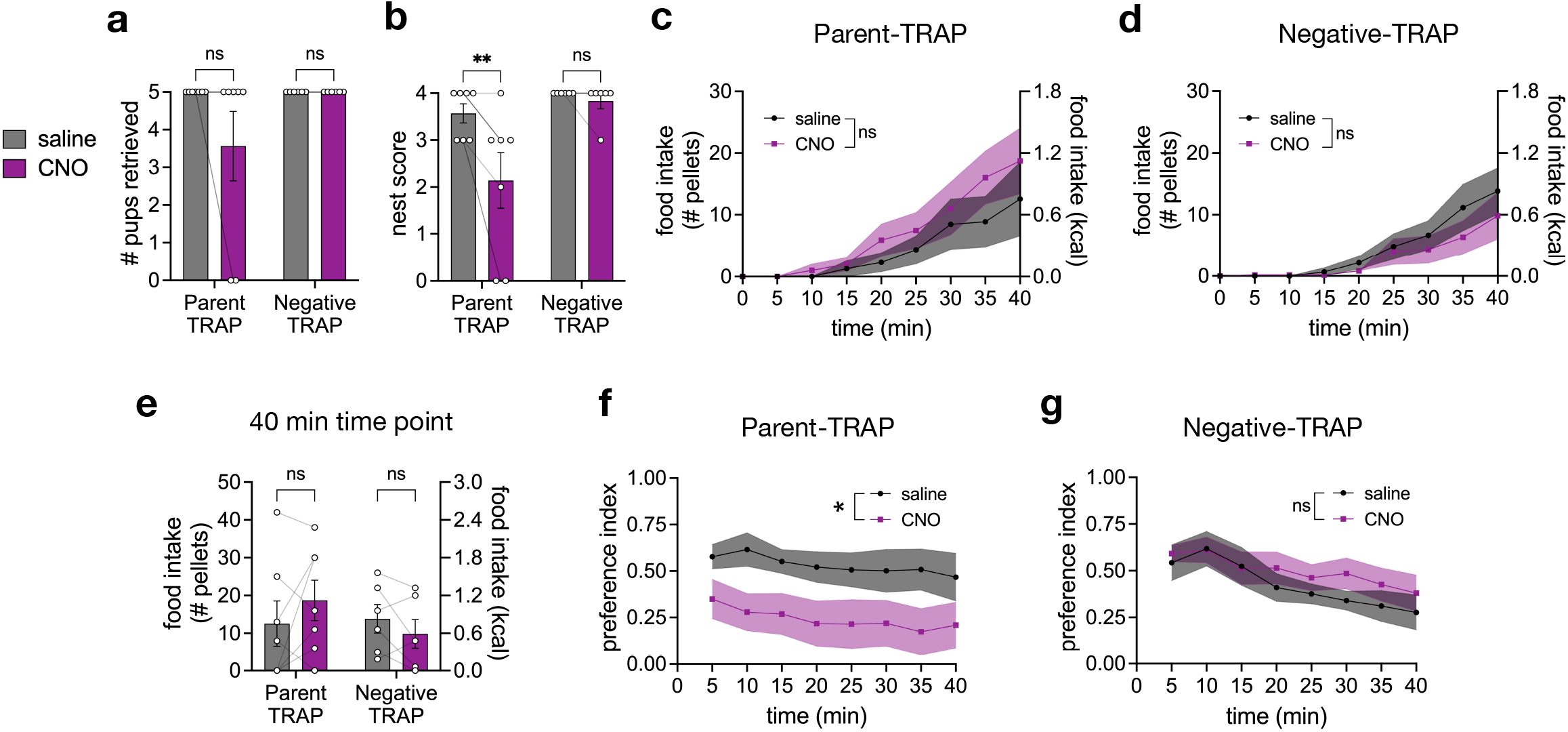
Inhibition of parenting neurons in MPOA reduces parenting in fed lactating mothers. **a-g**,data from conflict assay showing the number of pups retrieved (**a**), nest score (**b**), food intake over time in Parent-TRAP (**c**) and Negative-TRAP (**d**) animals, total food intake at 40 min time point (**e**), preference index over time in Parent-TRAP (**f**) and Negative-TRAP (**g**) animals (*n* = 7 Parent-TRAP lactating mothers, *n* = 6 Negative TRAP lactating mothers). Bar and line graphs show mean ± s.e.m. Box plots show median, upper/lower quartiles, and upper/lower extremes. Data were analyzed using two-way RM-ANOVA with Šidák’s multiple comparisons test. Significance symbols in **c, d, f, g** show the treatment effect of CNO. \**P* < 0.05, \*\**P* < 0.01, \*\*\**P* < 0.001, \*\*\*\**P* < 0.0001. ns, non-significant. See Supplementary Table 1 for detailed statistical information.

**Extended Data Fig. 8.**
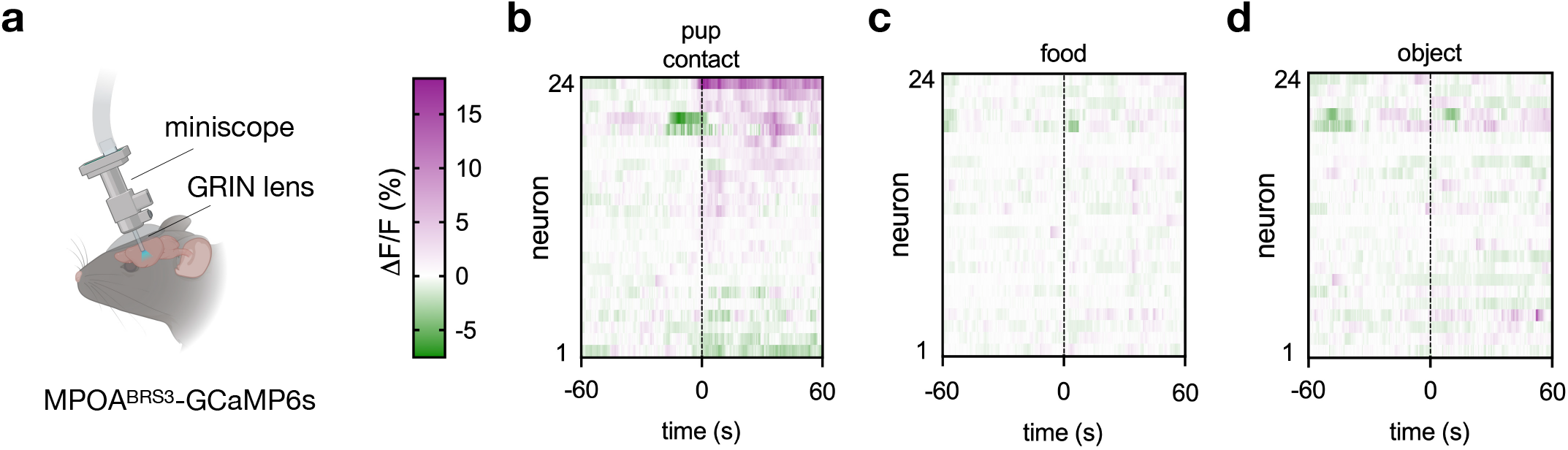
MPOA^BRS3^ neurons are responsive to pups. **a**, schematic of calcium imaging of MPOA^BRS3^-GCaMP6s using a miniscope; created with BioRender. **b-d**, calcium dynamics traces of individual MPOA^BRS3^ neurons in response to a pup that was retrieved (**b**), food (**c**), or an object (**d**) (*n* = 1 fed virgin female).

