## Supplementary Tables for "A Hypothalamic Circuit that Modulates Feeding and Parenting Behaviors"

### Supplementary Table 1: Statistical Information

| Figure | Statistical test | degrees of freedom (df), *t*-values, *F*-values F (DFn, DFd), *P* values, Pearson's r, Chi-square |
|---|---|---|
| **1b** | simple linear regression | Pearson's r = 0.8969<br>R squared = 0.8045<br>slope = 0.5496<br>slope $P < 0.0001$ |
| **1i** | unpaired t-test (two-tailed) | <u>firing rate individual neurons</u><br>t = 3.852<br>df = 180<br>$P = 0.0002$<br><br><u>mean firing rate per mouse</u><br>t = 2.438<br>df = 10<br>$P = 0.0350$ |
| **1k** | 2-way RM-ANOVA<br><br><br><br>Fisher's LSD test | hunger x lactation states: df = 1, F (1, 9) = 2.431, $P = 0.1534$<br>hunger state: df = 1, F (1, 9) = 48.15, $P < 0.0001$<br>lactation state: df = 1, F (1, 9) = 2.404, $P = 0.1555$<br><br>fed virgin vs. fed lactating: $P = 0.0461$<br>fasted virgin vs. fasted lactating: $P = 0.6351$<br>fed virgin vs. fasted virgin: $P = 0.0003$<br>fed lactating vs. fasted lactating: $P = 0.0032$ |
| **3h** | 2-way RM-ANOVA<br><br><br><br><br><br>Šídák's multiple comparisons test | lactation state x experimental condition: df = 2, F (2, 42) = 13.59, $P < 0.0001$<br>lactation state: df = 1, F (1, 21) = 32.88, $P < 0.0001$<br>experimental condition: df = 2, F (1.463, 30.73) = 116.7, $P < 0.0001$<br><br><u>virgin</u><br>fed (with pups) vs. fasted (with pups): $P < 0.0001$<br>fed (with pups) vs. fasted (no pups): $P = 0.0002$<br>fasted (with pups) vs. fasted (no pups): $P = 0.2045$<br><br><u>lactating</u><br>fed (with pups) vs. fasted (with pups): $P < 0.0001$<br>fed (with pups) vs. fasted (no pups): $P < 0.0001$<br>fasted (with pups) vs. fasted (no pups): $P = 0.0003$ |
| **3i** | 2-way RM-ANOVA<br><br><br><br>Šídák's multiple comparisons test | lactation x hunger states: df = 1, F (1, 21) = 9.995, $P = 0.0047$<br>lactation state: df = 1, F (1, 21) = 7.418, $P = 0.0127$<br>hunger state: df = 1, F (1, 21) = 9.995, $P = 0.0047$<br><br>fed virgin vs. fasted virgin: $P = 0.0008$<br>fed lactating vs. fasted lactating: $P > 0.9999$ |
| **3j** | 2-way RM-ANOVA<br><br><br><br>Šídák's multiple comparisons test | pup x hunger state: df = 1, F (1, 12) = 5.208, $P = 0.0415$<br>pup: df = 1, F (1, 12) = 36.60, $P < 0.0001$<br>hunger state: df = 1, F (1, 12) = 4.283, $P = 0.0607$<br><br>first pup, fed vs. fasted: $P = 0.9940$<br>last pup, fed vs. fasted: $P = 0.0120$ |

| Figure | Statistical test | degrees of freedom (df), *t*-values, *F*-values F (DFn, DFd), *P* values, Pearson's r, Chi-square |
|---|---|---|
| **3k** | 2-way RM-ANOVA<br><br><br><br><br><br>Šídák's multiple comparisons test | lactation state x experimental condition: df = 2, F (2, 42) = 0.6610, *P* = 0.5216<br>lactation state: df = 1, F (1, 21) = 0.6718, *P* = 0.4216<br>experimental condition: df = 2, F (1.303, 27.36) = 573.8, *P* < 0.0001<br><br><u>virgin</u><br>fed (with pups) vs. fasted (with pups): *P* = 0.2606<br>fed (with pups) vs. fasted (no pups): *P* < 0.0001<br>fasted (with pups) vs. fasted (no pups): *P* < 0.0001<br><br><u>lactating</u><br>fed (with pups) vs. fasted (with pups): *P* = 0.2266<br>fed (with pups) vs. fasted (no pups): *P* < 0.0001<br>fasted (with pups) vs. fasted (no pups): *P* < 0.0001 |
| **3l** | 2-way RM-ANOVA<br><br><br><br><br><br>Šídák's multiple comparisons test | lactation state x experimental condition: df = 2, F (2, 42) = 1.472, *P* = 0.2411<br>lactation state: df = 1, F (1, 21) = 10.01, *P* = 0.0047<br>experimental condition: df = 2, F (1.901, 39.92) = 132.2, *P* < 0.0001<br><br><u>virgin</u><br>fed (with pups) vs. fasted (with pups): *P* = 0.0046<br>fed (with pups) vs. fasted (no pups): *P* < 0.0001<br>fasted (with pups) vs. fasted (no pups): *P* < 0.0001<br><br><u>lactating</u><br>fed (with pups) vs. fasted (with pups): *P* < 0.0001<br>fed (with pups) vs. fasted (no pups): *P* < 0.0001<br>fasted (with pups) vs. fasted (no pups): *P* = 0.0005 |
| **3m** | simple linear regression | <u>virgin</u><br>Pearson's r = 0.6930<br>R squared = 0.4803<br>slope = -0.3382<br>slope *P* = 0.0263<br><br><u>lactating</u><br>Pearson's r = 0.6298<br>R squared = 0.3966<br>slope = -0.8520<br>slope *P* < 0.0211 |

| Figure | Statistical test | degrees of freedom (df), *t*-values, *F*-values F (DFn, DFd), *P* values, Pearson's r, Chi-square |
|---|---|---|
| **4c** | 2-way RM-ANOVA<br><br><br><br>Šídák's multiple comparisons test | light intensity x paradigm: df = 3, F (2.342, 21.08) = 0.5339, *P* = 0.6215<br>light intensity: df = 1, F (1.000, 9.000) = 2.808, *P* = 0.1281<br>paradigm: df = 3, F (1.823, 16.41) = 115.3, *P* < 0.0001<br><br>10 mW<br>no stim vs. pre-stim: *P* < 0.0001<br>no stim vs. co-stim: *P* < 0.0001<br>no stim vs. pre- & co-stim: *P* < 0.0001<br>pre-stim vs. co-stim: *P* = 0.0380<br>pre-stim vs. pre- & co-stim: *P* = 0.0033<br>co-stim vs. pre- & co-stim: *P* = 0.0022<br><br>1 mW<br>no stim vs. pre-stim: *P* = 0.0007<br>no stim vs. co-stim: *P* = 0.0002<br>no stim vs. pre- & co-stim: *P* < 0.0001<br>pre-stim vs. co-stim: *P* = 0.0295<br>pre-stim vs. pre- & co-stim: *P* = 0.0015<br>co-stim vs. pre- & co-stim: *P* = 0.2619 |
| **4d** | Mixed-effects model<br><br><br><br>Šídák's multiple comparisons test | lactation state x experimental condition: F (1.249, 9.367) = 8.368, *P* = 0.0137<br>lactation state: F (1.000, 9.000) = 21.61, *P* = 0.0012<br>experimental condition: F (1.282, 11.54) = 51.82, *P* < 0.0001<br><br>virgin<br>fed (no stim) vs. fed (stim): *P* < 0.0001<br>fed (no stim) vs. fasted (no stim): *P* = 0.0001<br>fed (stim) vs. fasted (no stim): *P* = 0.0031<br><br>lactating<br>fed (no stim) vs. fed (stim): *P* = 0.0009<br>fed (no stim) vs. fasted (no stim): *P* < 0.0001<br>fed (stim) vs. fasted (no stim): *P* = 0.9624 |
| **4e** | Mixed-effects model<br><br><br><br>Šídák's multiple comparisons test | lactation state x experimental condition: F (1.396, 10.47) = 0.4315, *P* = 0.5911<br>lactation state: F (1.000, 9.000) = 0.01757, *P* = 0.8975<br>experimental condition: F (1.082, 9.740) = 17.66, *P* = 0.0017<br><br>virgin<br>fed (no stim) vs. fed (stim): *P* = 0.0091<br>fed (no stim) vs. fasted (no stim): *P* = 0.2606<br>fed (stim) vs. fasted (no stim): *P* = 0.0228<br><br>lactating<br>fed (no stim) vs. fed (stim): *P* = 0.0118<br>fed (no stim) vs. fasted (no stim): *P* = 0.2226<br>fed (stim) vs. fasted (no stim): *P* = 0.0131 |

| Figure | Statistical test | degrees of freedom (df), *t*-values, *F*-values F (DFn, DFd), *P* values, Pearson's r, Chi-square |
|---|---|---|
| **4f** | Mixed-effects model<br><br><br><br><br>Šídák's multiple comparisons test | lactation state x experimental condition: F (1.991, 14.93) = 1.436, *P* = 0.2689<br>lactation state: F (1.000, 9.000) = 6.729, *P* = 0.0290<br>experimental condition: F (1.196, 10.77) = 27.56, *P* = 0.0002<br><br><u>virgin</u><br>fed (no stim) vs. fed (stim): *P* = 0.0118<br>fed (no stim) vs. fasted (no stim): *P* = 0.2242<br>fed (stim) vs. fasted (no stim): *P* = 0.0153<br><br><u>lactating</u><br>fed (no stim) vs. fed (stim): *P* = 0.0006<br>fed (no stim) vs. fasted (no stim): *P* = 0.1833<br>fed (stim) vs. fasted (no stim): *P* = 0.0004 |
| **4g** | Mixed-effects model<br><br><br><br><br>Tukey's multiple comparisons test | lactation state x experimental condition: F (1.419, 10.64) = 2.216, *P* = 0.1632<br>lactation state: F (1.000, 9.000) = 13.94, *P* = 0.0047<br>experimental condition: F (1.965, 17.68) = 22.32, *P* < 0.0001<br><br><u>virgin</u><br>fed (no stim) vs. fed (stim): *P* = 0.0095<br>fed (no stim) vs. fasted (no stim): *P* = 0.1422<br>fed (stim) vs. fasted (no stim): *P* = 0.0163<br><br><u>lactating</u><br>fed (no stim) vs. fed (stim): *P* = 0.0031<br>fed (no stim) vs. fasted (no stim): *P* = 0.0968<br>fed (stim) vs. fasted (no stim): *P* = 0.03814 |
| **5b** | 2-way RM-ANOVA<br><br><br><br>Šídák's multiple comparisons test | TRAP x drug: df = 1, F (1, 11) = 41.05, *P* <0.0001<br>TRAP: df = 1, F (1, 11) = 41.05, *P* <0.0001<br>drug: df = 1, F (1, 11) = 41.05, *P* <0.0001<br><br>Parent-TRAP, saline vs. CNO: *P* < 0.0001<br>Negative-TRAP, saline vs. CNO: *P* > 0.9999 |
| **5c** | 2-way RM-ANOVA<br><br><br><br>Šídák's multiple comparisons test | TRAP x drug: df = 1, F (1, 11) = 27.64, *P* = 0.0003<br>TRAP: df = 1, F (1, 11) = 48.81, *P* < 0.0001<br>drug: df = 1, F (1, 11) = 44.17, *P* <0.0001<br><br>Parent-TRAP, saline vs. CNO: *P* < 0.0001<br>Negative-TRAP, saline vs. CNO: *P* = 0.5960 |

| Figure | Statistical test | degrees of freedom (df), *t*-values, *F*-values F (DFn, DFd), *P* values, Pearson's r, Chi-square |
|---|---|---|
| **5d** | 2-way RM-ANOVA<br><br><br><br>Šídák's multiple comparisons test | time x drug: df = 6, F (6, 36) = 2.712, *P* = 0.0282<br>time: df = 6, F (6, 36) = 233.8, *P* < 0.0001<br>drug: df = 1, F (1, 6) = 2.006, *P* = 0.2064<br><br><u>saline vs. CNO</u><br>5 min: *P* = 0.1264<br>10 min: *P* = 0.0024<br>15 min: *P* = 0.8838<br>20 min: *P* = 0.9996<br>25 min: *P* > 0.9999<br>30 min: *P* = 0.9685<br>35 min: *P* > 0.9999 |
| **5e** | 2-way RM-ANOVA | time x drug: df = 8, F (8, 40) = 0.7731, *P* = 0.6283<br>time: df = 8, df = 8, F (8, 40) = 200.7, *P* < 0.0001<br>drug: df = 1, F (1, 5) = 1.164, *P* = 0.3300 |
| **5f** | 2-way RM-ANOVA<br><br><br><br>Šídák's multiple comparisons test | TRAP x drug: df = 1, F (1, 12) = 3.089, *P* = 0.1043<br>TRAP: df = 1, F (1, 12) = 1.379, *P* = 0.2630<br>drug: df = 1, F (1, 11) = 44.17, *P* = 0.0719<br><br>Parent-TRAP, saline vs. CNO: *P* = 0.0290<br>Negative-TRAP, saline vs. CNO: *P* = 0.9876 |
| **5g** | 2-way RM-ANOVA<br><br><br><br>Šídák's multiple comparisons test | TRAP x drug: df = 1, F (1, 12) = 0.8103, *P* = 0.3857<br>TRAP: df = 1, F (1, 12) = 2.394, *P* = 0.1477<br>drug: df = 1, F (1, 12) = 3.782, *P* = 0.0756<br><br>Parent-TRAP, saline vs. CNO: *P* = 0.0985<br>Negative-TRAP, saline vs. CNO: *P* = 0.7528 |
| **5h** | 2-way RM-ANOVA | time x drug: df = 7, F (1.444, 8.666) = 4.576, *P* = 0.0526<br>time: df = 7, F (1.808, 10.85) = 9.912, *P* = 0.0041<br>drug: df = 1, F (1.000, 6.000) = 59.69, *P* = 0.0002 |
| **5i** | 2-way RM-ANOVA | time x drug: df = 7, F (7, 35) = 0.9033, *P* = 0.5150<br>time: df = 7, df = 8, F (7, 35) = 2.911, *P* = 0.0165<br>drug: df = 1, F (1, 5) = 1.857, *P* = 0.2311 |
| **5j** | 2-way RM-ANOVA<br><br><br><br>Šídák's multiple comparisons test | TRAP x drug: df = 1, F (1, 11) = 26.17, *P* = 0.0003<br>TRAP: df = 1, F (1, 11) = 0.8599, *P* = 0.3737<br>drug: df = 1, F (1, 11) = 21.10, *P* = 0.0008<br><br>Parent-TRAP, saline vs. CNO: *P* < 0.0001<br>Negative-TRAP, saline vs. CNO: *P* = 0.9264 |
| **5k** | 2-way RM-ANOVA<br><br><br><br>Šídák's multiple comparisons test | TRAP x drug: df = 1, F (1, 11) = 3.223, *P* = 0.1001<br>TRAP: df = 1, F (1, 11) = 0.1748, *P* = 0.6839<br>drug: df = 1, F (1, 11) = 10.33, *P* = 0.0083<br><br>Parent-TRAP, saline vs. CNO: *P* = 0.0072<br>Negative-TRAP, saline vs. CNO: *P* = 0.5835 |

| Figure | Statistical test | degrees of freedom (df), *t*-values, *F*-values F (DFn, DFd), *P* values, Pearson's r, Chi-square |
|---|---|---|
| **6a** | 2-way RM-ANOVA<br><br><br><br>Fisher's LSD test | hunger x drug: df = 1, F (1.000, 7.000) = 0.08046, *P* = 0.7849<br>hunger: df = 1, F (1.000, 7.000) = 0.9726, *P* = 0.3569<br>drug: df = 1, F (1.000, 7.000) = 23.84, *P* = 0.0018<br><br>fed, saline vs. CNO: *P* = 0.0071<br>fasted, saline vs. CNO: *P* = 0.0331<br>saline, fed vs. fasted: *P* = 0.1114<br>CNO, fed vs. fasted: *P* = 0.7567 |
| **6b** | 2-way RM-ANOVA<br><br><br><br>Fisher's LSD test | hunger x drug: df = 1, F (1.000, 7.000) = 5.600, *P* = 0.0499<br>hunger: df = 1, F (1.000, 7.000) = 1.800, *P* = 0.2216<br>drug: df = 1, F (1.000, 7.000) = 46.07, *P* = 0.0003<br><br>fed, saline vs. CNO: *P* = 0.0061<br>fasted, saline vs. CNO: *P* < 0.0001<br>saline, fed vs. fasted: *P* = 0.5983<br>CNO, fed vs. fasted: *P* = 0.0875 |
| **6c** | 2-way RM-ANOVA<br><br><br><br>Fisher's LSD test | hunger x drug: df = 1, F (1.000, 7.000) = 6.724, *P* = 0.0358<br>hunger: df = 1, F (1.000, 7.000) = 3.438, *P* = 0.1061<br>drug: df = 1, F (1.000, 7.000) = 13.45, *P* = 0.0080<br><br>fed, saline vs. CNO: *P* = 0.0052<br>fasted, saline vs. CNO: *P* = 0.7854<br>saline, fed vs. fasted: *P* = 0.0136<br>CNO, fed vs. fasted: *P* = 0.7412 |
| **6d** | 2-way RM-ANOVA<br><br><br><br>Fisher's LSD test | hunger x drug: df = 1, F (1.000, 7.000) = 0.7422, *P* = 0.4175<br>hunger: df = 1, F (1.000, 7.000) = 70.91, *P* < 0.0001<br>drug: df = 1, F (1.000, 7.000) = 2.156, *P* = 0.1855<br><br>fed, saline vs. CNO: *P* = 0.0263<br>fasted, saline vs. CNO: *P* = 0.2861<br>saline, fed vs. fasted: *P* < 0.0001<br>CNO, fed vs. fasted: *P* = 0.0017 |
| **6e** | 2-way RM-ANOVA | time x drug: df = 4, F (1.553, 9.319) = 10.14, *P* = 0.0063<br>time: df = 4, F (1.715, 10.29) = 19.42, *P* = 0.0004<br>drug: df = 1, F (1.000, 6.000) = 14.42, *P* = 0.0090 |
| **6f** | Chi-square test | Chi-square = 7.778<br>df = 2<br>*P* = 0.0205 |
| **6g** | 2-way RM-ANOVA | time x drug: df = 4, F (2.143, 10.71) = 1.092, *P* = 0.3746<br>time: df = 4, F (1.455, 7.275) = 109.7, *P* < 0.0001<br>drug: df = 1, F (1.000, 5.000) = 1.367, *P* = 0.2951 |
| **6k** | Wilcoxon matched-pairs signed rank test (two-tailed) | *P* = 0.0312 |
| **6m** | Wilcoxon matched-pairs signed rank test (two-tailed) | *P* = 0.0312 |

| Figure | Statistical test | degrees of freedom (df), *t*-values, *F*-values F (DFn, DFd), *P* values, Pearson's r, Chi-square |
|---|---|---|
| **Ext. 1a** | 2-way RM-ANOVA<br><br><br>Šídák's multiple comparisons test | time x lactation df = 3, F (3, 60) = 132.8, *P* < 0.0001<br>time df = 3, F (1.052, 21.04) = 121.0, *P* < 0.0001<br>lactation df = 1, F (1, 20) = 114.3, *P* < 0.0001<br><br>control vs. lactating<br>baseline *P* > 0.9999<br>pregnancy *P* < 0.0001<br>lactation *P* < 0.0001<br>post-weaning *P* < 0.0001 |
| **Ext. 1e** | 2-way RM-ANOVA<br><br><br>Šídák's multiple comparisons test | time x lactation df = 3, F (3, 27) = 119.5, *P* < 0.0001<br>time df = 3, F (2.036, 18.32) = 98.55, *P* < 0.0001<br>lactation df = 1, F (1, 9) = 40.32, *P* = 0.0001<br><br>control vs. lactating<br>baseline *P* = 0.6266<br>pregnancy *P* = 0.4433<br>lactation *P* = 0.0001<br>post-weaning *P* = 0.0351 |
| **Ext. 1g** | 2-way RM-ANOVA<br><br><br>Šídák's multiple comparisons test | time x lactation df = 3, F (3, 27) = 66.12, *P* < 0.0001<br>time df = 3, F (1.448, 13.03) = 54.85, *P* < 0.0001<br>lactation df = 1, F (1, 9) = 12.65, *P* = 0.0062<br><br>control vs. lactating<br>baseline *P* = 0.9985<br>pregnancy *P* = 0.9519<br>lactation *P* = 0.0086<br>post-weaning *P* = 0.8097 |
| **Ext. 5c** | paired t-test (two-tailed) | t = 1.431<br>df = 5<br>*P* = 0.2117 |
| **Ext. 5d** | 1-way RM-ANOVA | F = 50.62<br>*P* = 0.0008<br><br>fed vs. fed (stim): df = 5, *P* = 0.6700<br>fed vs. fasted: df = 5, *P* = 0.0024<br>fed (stim) vs. fasted: df = 5, *P* = 0.0015 |
| **Ext. 5e** | Friedman test<br><br><br>Dunn's multiple comparisons test | Friedman statistic = 6.000<br>*P* = 0.1111<br><br>fed vs. fed (stim): *P* > 0.9999<br>fed vs. fasted: *P* = 0.5818<br>fed (stim) vs. fasted: *P* = 0.5818 |
| **Ext. 5f** | Friedman test<br><br><br>Dunn's multiple comparisons test | Friedman statistic = 2.000<br>*P* > 0.9999<br><br>fed vs. fed (stim): *P* > 0.9999<br>fed vs. fasted: *P P* > 0.9999<br>fed (stim) vs. fasted: *P P* > 0.9999 |

| Figure | Statistical test | degrees of freedom (df), *t*-values, *F*-values F (DFn, DFd), *P* values, Pearson's r, Chi-square |
|---|---|---|
| **Ext. 5g** | 1-way RM-ANOVA | F = 50.62<br>*P* = 0.0008<br><br>fed vs. fed (stim): df = 5, *P* = 0.6700<br>fed vs. fasted: df = 5, *P* = 0.0024<br>fed (stim) vs. fasted: df = 5, *P* = 0.0015 |
| **Ext. 6a**<br><br>**food intake** | 2-way RM-ANOVA<br><br><br><br>Fisher's LSD test | hunger x drug: df = 1, F (1.000, 7.000) = 31.59, *P* = 0.0008<br>hunger: df = 1, F (1.000, 7.000) = 105.2, *P* < 0.0001<br>drug: df = 1, F (1.000, 7.000) = 29.21, *P* = 0.0010<br><br>fed, saline vs. CNO: *P* = 0.4015<br>fasted, saline vs. CNO: *P* = 0.0007<br>saline, fed vs. fasted: *P* < 0.0001<br>CNO, fed vs. fasted: *P* = 0.0016 |
| **Ext. 6a**<br><br>**pup retrieval** | 2-way RM-ANOVA<br><br><br><br>Fisher's LSD test | hunger x drug: df = 1, F (1.000, 7.000) = 0.2195, *P* = 0.6537<br>hunger: df = 1, F (1.000, 7.000) = 1.127, *P* = 0.3236<br>drug: df = 1, F (1.000, 7.000) = 47.54, *P* = 0.0002<br><br>fed, saline vs. CNO: *P* = 0.0013<br>fasted, saline vs. CNO: *P* = 0.0011<br>saline, fed vs. fasted: *P* = 0.3645<br>CNO, fed vs. fasted: *P* = 0.5040 |
| **Ext. 6a**<br><br>**nest score** | 2-way RM-ANOVA<br><br><br><br>Fisher's LSD test | hunger x drug: df = 1, F (1.000, 7.000) = 0.1273, *P* = 0.7318<br>hunger: df = 1, F (1.000, 7.000) = 1.842, *P* = 0.2168<br>drug: df = 1, F (1.000, 7.000) = 127.7, *P* < 0.0001<br><br>fed, saline vs. CNO: *P* < 0.0001<br>fasted, saline vs. CNO: *P* = 0.0001<br>saline, fed vs. fasted: *P* = 0.1970<br>CNO, fed vs. fasted: *P* = 0.4512 |
| **Ext. 6a**<br><br>**preference index** | 2-way RM-ANOVA<br><br><br><br>Fisher's LSD test | hunger x drug: df = 1, F (1.000, 7.000) = 9.022, *P* = 0.0198<br>hunger: df = 1, F (1.000, 7.000) = 0.2426, *P* = 0.6374<br>drug: df = 1, F (1.000, 7.000) = 18.05, *P* = 0.0038<br><br>fed, saline vs. CNO: *P* = 0.0019<br>fasted, saline vs. CNO: *P* = 0.5070<br>saline, fed vs. fasted: *P* = 0.0752<br>CNO, fed vs. fasted: *P* = 0.1226 |
| **Ext. 6b**<br><br>**food intake** | 2-way RM-ANOVA<br><br><br><br>Fisher's LSD test | hunger x drug: df = 1, F (1.000, 9.000) = 0.09484, *P* = 0.7651<br>hunger: df = 1, F (1.000, 9.000) = 60.28, *P* < 0.0001<br>drug: df = 1, F (1.000, 9.000) = 0.01197, *P* = 0.9153<br><br>fed, saline vs. CNO: *P* = 0.2569<br>fasted, saline vs. CNO: *P* = 0.8364<br>saline, fed vs. fasted: *P* = 0.0005<br>CNO, fed vs. fasted: *P* = 0.0006 |

| Figure | Statistical test | degrees of freedom (df), *t*-values, *F*-values F (DFn, DFd), *P* values, Pearson's r, Chi-square |
|---|---|---|
| **Ext. 6b**<br><br>**pup retrieval** | 2-way RM-ANOVA<br><br><br>Fisher's LSD test | hunger x drug: df = 1, F (1.000, 9.000) = 8.265, *P* = 0.0183<br>hunger: df = 1, F (1.000, 9.000) = 13.96, *P* = 0.0047<br>drug: df = 1, F (1.000, 9.000) = 7.759, *P* = 0.0212<br><br>fed, saline vs. CNO: *P* = 0.1382<br>fasted, saline vs. CNO: *P* = 0.0150<br>saline, fed vs. fasted: *P* = 0.1679<br>CNO, fed vs. fasted: *P* = 0.0044 |
| **Ext. 6b**<br><br>**nest score** | 2-way RM-ANOVA<br><br><br>Fisher's LSD test | hunger x drug: df = 1, F (1.000, 9.000) = 4.091, *P* = 0.0738<br>hunger: df = 1, F (1.000, 9.000) = 7.230, *P* = 0.0248<br>drug: df = 1, F (1.000, 9.000) = 22.22, *P* = 0.0011<br><br>fed, saline vs. CNO: *P* = 0.0051<br>fasted, saline vs. CNO: *P* = 0.0063<br>saline, fed vs. fasted: *P* = 0.1679<br>CNO, fed vs. fasted: *P* = 0.0368 |
| **Ext. 6b**<br><br>**preference index** | 2-way RM-ANOVA<br><br><br>Fisher's LSD test | hunger x drug: df = 1, F (1.000, 9.000) = 0.02990, *P* = 0.8665<br>hunger: df = 1, F (1.000, 9.000) = 132.0, *P* < 0.0001<br>drug: df = 1, F (1.000, 9.000) = 0.9487, *P* = 0.3555<br><br>fed, saline vs. CNO: *P* = 0.1944<br>fasted, saline vs. CNO: *P* = 0.5442<br>saline, fed vs. fasted: *P* = 0.0002<br>CNO, fed vs. fasted: *P* = 0.0002 |
| **Ext. 6c**<br><br>**food intake** | 2-way RM-ANOVA<br><br><br>Fisher's LSD test | hunger x drug: df = 1, F (1.000, 5.000) = 0.4634, *P* = 0.5263<br>hunger: df = 1, F (1.000, 5.000) = 18.97, *P* = 0.0073<br>drug: df = 1, F (1.000, 5.000) = 0.3296, *P* = 0.5908<br><br>fed, saline vs. CNO: *P* = 0.3632<br>fasted, saline vs. CNO: *P* = 0.5563<br>saline, fed vs. fasted: *P* = 0.0020<br>CNO, fed vs. fasted: *P* = 0.0224 |
| **Ext. 6c**<br><br>**pup retrieval** | 2-way RM-ANOVA<br><br><br>Fisher's LSD test | hunger x drug: df = 1, F (1.000, 5.000) = 2.143, *P* = 0.2031<br>hunger: df = 1, F (1.000, 5.000) = 2.500, *P* = 0.1747<br>drug: df = 1, F (1.000, 5.000) = 2.143, *P* = 0.2031<br><br>fed, saline vs. CNO: no difference<br>fasted, saline vs. CNO: *P* = 0.2031<br>saline, fed vs. fasted: *P* = 0.1801<br>CNO, fed vs. fasted: *P* = 0.3632 |
| **Ext. 6c**<br><br>**nest score** | 2-way RM-ANOVA<br><br><br>Fisher's LSD test | hunger x drug: df = 1, F (1.000, 5.000) = 1.000, *P* = 0.3632<br>hunger: df = 1, no effect<br>drug: df = 1, no effect<br><br>fed, saline vs. CNO: *P* = 0.3632<br>fasted, saline vs. CNO: *P* = 0.3632<br>saline, fed vs. fasted: *P* = 0.3632<br>CNO, fed vs. fasted: *P* = 0.3632 |

| Figure | Statistical test | degrees of freedom (df), *t*-values, *F*-values F (DFn, DFd), *P* values, Pearson's r, Chi-square |
|---|---|---|
| **Ext. 6c**<br><br>**preference index** | 2-way RM-ANOVA<br><br><br><br>Fisher's LSD test | hunger x drug: df = 1, F (1.000, 5.000) = 0.1595, *P* = 0.7061<br>hunger: df = 1, F (1.000, 5.000) = 11.20, *P* = 0.0204<br>drug: df = 1, F (1.000, 5.000) = 1.555, *P* = 0.2676<br><br>fed, saline vs. CNO: *P* = 0.2640<br>fasted, saline vs. CNO: *P* = 0.6878<br>saline, fed vs. fasted: *P* = 0.0887<br>CNO, fed vs. fasted: *P* = 0.0025 |
| **Ext. 6d**<br><br>**food intake** | 2-way RM-ANOVA<br><br><br><br>Fisher's LSD test | hunger x drug: df = 1, F (1.000, 5.000) = 1.121, *P* = 0.3382<br>hunger: df = 1, F (1.000, 5.000) = 43.88, *P* = 0.0012<br>drug: df = 1, F (1.000, 5.000) = 0.2123, *P* = 0.6643<br><br>fed, saline vs. CNO: *P* = 0.2234<br>fasted, saline vs. CNO: *P* = 0.8890<br>saline, fed vs. fasted: *P* = 0.0035<br>CNO, fed vs. fasted: *P* = 0.0018 |
| **Ext. 6d**<br><br>**pup retrieval** | N/A | N/A (values across all conditions are identical) |
| **Ext. 6d**<br><br>**nest score** | 2-way RM-ANOVA<br><br><br><br>Fisher's LSD test | hunger x drug: df = 1, F (1.000, 5.000) = 5.000, *P* = 0.0756<br>hunger: df = 1, F (1.000, 5.000) = 0.1220, *P* = 0.7412<br>drug: df = 1, F (1.000, 5.000) = 0.2941, *P* = 0.6109<br><br>fed, saline vs. CNO: *P* = 0.1747<br>fasted, saline vs. CNO: *P* = 0.3632<br>saline, fed vs. fasted: *P* = 0.6109<br>CNO, fed vs. fasted: *P* = 0.1747 |
| **Ext. 6d**<br><br>**preference index** | 2-way RM-ANOVA<br><br><br><br>Fisher's LSD test | hunger x drug: df = 1, F (1.000, 5.000) = 0.07216, *P* = 0.7990<br>hunger: df = 1, F (1.000, 5.000) = 21.38, *P* = 0.0057<br>drug: df = 1, F (1.000, 5.000) = 0.0003487, *P* = 0.9858<br><br>fed, saline vs. CNO: *P* = 0.8994<br>fasted, saline vs. CNO: *P* = 0.9433<br>saline, fed vs. fasted: *P* = 0.0233<br>CNO, fed vs. fasted: *P* = 0.0012 |
| **Ext. 7a** | 2-way RM-ANOVA<br><br><br><br>Šídák's multiple comparisons test | TRAP x drug: df = 1, F (1, 11) = 2.031, *P* = 0.1819<br>TRAP: df = 1, F (1, 11) = 2.031, *P* = 0.1819<br>drug: df = 1, F (1, 11) = 2.031, *P* = 0.1819<br><br>Parent-TRAP, saline vs. CNO: *P* = 0.1161<br>Negative-TRAP, saline vs. CNO: *P* > 0.9999 |
| **Ext. 7b** | 2-way RM-ANOVA<br><br><br><br>Šídák's multiple comparisons test | TRAP x drug: df = 1, F (1, 11) = 6.621, *P* = 0.0259<br>TRAP: df = 1, F (1, 11) = 6.077, *P* = 0.0314<br>drug: df = 1, F (1, 11) = 10.58, *P* = 0.0077<br><br>Parent-TRAP, saline vs. CNO: *P* = 0.0026<br>Negative-TRAP, saline vs. CNO: *P* = 0.8791 |
| **Ext. 7c** | 2-way RM-ANOVA | time x drug: df = 8, F (1.451, 8.705) = 1.252, *P* = 0.3161<br>time: df = 8, F (1.212, 7.272) = 9.495, *P* = 0.0145<br>drug: df = 1, F (1.000, 6.000) = 1.194, *P* = 0.3164 |

| Figure | Statistical test | degrees of freedom (df), *t*-values, *F*-values F (DFn, DFd), *P* values, Pearson's r, Chi-square |
|---|---|---|
| **Ext. 7d** | 2-way RM-ANOVA | time x drug: df = 8, F (2.367, 11.84) = 1.695, *P* = 0.2244<br>time: df = 8, F (1.203, 6.016) = 9.242, *P* = 0.0203<br>drug: df = 1, F (1.000, 5.000) = 2.082, *P* = 0.2086 |
| **Ext. 7e** | 2-way RM-ANOVA<br><br><br>Šídák's multiple comparisons test | TRAP x drug: df = 1, F (1, 11) = 2.398, *P* = 0.1498<br>TRAP: df = 1, F (1, 11) = 0.3665, *P* = 0.5572<br>drug: df = 1, F (1, 11) = 0.1070, *P* = 0.7497<br><br>Parent-TRAP, saline vs. CNO: *P* = 0.3518<br>Negative-TRAP, saline vs. CNO: *P* = 0.6671 |
| **Ext. 7f** | 2-way RM-ANOVA | time x drug: df = 7, F (2.179, 13.08) = 0.4135, *P* = 0.6863<br>time: df = 7, F (1.204, 7.227) = 0.7302, *P* = 0.4461<br>drug: df = 1, F (2.179, 13.08) = 0.4135, *P* = 0.0391 |
| **Ext. 7g** | 2-way RM-ANOVA | time x drug: df = 7, F (7, 35) = 0.9033, *P* = 0.5150<br>time: df = 7, F (7, 35) = 2.911, *P* = 0.0165<br>drug: df = 1, F (1, 5) = 1.857, *P* = 0.2311 |

**Supplementary Table 2:**

Differentially expressed genes in *Agrp*+ ARC neuronal cluster. Positive values in the “avg_log2FC” columns indicate upregulation in the fasted and lactating states.

| gene | avg_log2FC_fasted | p_val_adj_fasted | avg_log2FC_lactating | p_val_adj_lactating |
|---|---|---|---|---|
| Vgf | 2.91181049 | 2.99E-180 | 0.44084309 | 1.01E-06 |
| Agrp | 1.44705604 | 1.69E-92 | 1.38557687 | 1.12E-102 |
| Npy | 1.39988235 | 1.30E-84 | 1.23823864 | 9.81E-105 |
| Lepr | 1.31048548 | 1.17E-94 | 0.37054165 | 1.32E-10 |
| Rmst | 1.25634306 | 8.67E-75 | 0.49550606 | 1.02E-14 |
| 9630014M24Rik | 1.19464046 | 1.42E-101 | 0.38435819 | 9.43E-16 |
| Kcnc2 | 1.10515228 | 3.32E-58 | 0.44607897 | 7.81E-17 |
| Pex5l | 1.09574265 | 1.03E-60 | 0.31928216 | 1.28E-05 |
| Lrrc7 | 1.04826622 | 4.12E-58 | 0.47197013 | 1.04E-18 |
| Lancl3 | 0.99421541 | 2.73E-78 | 0.31629417 | 3.66E-13 |
| Acvr1c | 0.93664619 | 1.22E-42 | 0.61834722 | 7.33E-27 |
| Ghr | 0.9325291 | 6.06E-80 | 0.32504335 | 4.56E-13 |
| Hdac9 | 0.89622287 | 2.52E-28 | 0.5862532 | 1.93E-14 |
| 1500009L16Rik | 0.82690458 | 4.89E-48 | 0.25221041 | 6.80E-07 |
| Camk2d | 0.8160303 | 5.74E-51 | 0.31813305 | 7.17E-05 |
| Slc8a1 | 0.8094826 | 1.33E-33 | 0.32389502 | 3.94E-06 |
| Elmo1 | 0.74486697 | 2.51E-32 | 0.48991193 | 1.19E-22 |
| Cdh18 | 0.7371266 | 3.76E-28 | 0.6869603 | 5.80E-26 |
| Peg10 | 0.68102166 | 8.96E-14 | 0.26637774 | 1.65E-05 |
| Rasgef1b | 0.64368138 | 2.47E-21 | 0.47473239 | 5.58E-13 |
| Adcy2 | 0.6016727 | 7.63E-23 | 0.43042173 | 1.01E-09 |
| Pid1 | 0.59714633 | 2.32E-09 | 0.36648629 | 0.01183002 |
| Tbc1d4 | 0.58534006 | 1.47E-24 | 0.31206991 | 0.00022448 |
| Hcn1 | 0.56814321 | 5.12E-21 | 0.31152674 | 2.24E-05 |
| Pcp4 | 0.56327788 | 2.32E-17 | 0.67118689 | 5.76E-34 |

| gene | avg_log2FC_fasted | p_val_adj_fasted | avg_log2FC_lactating | p_val_adj_lactating |
|---|---|---|---|---|
| Grm8 | 0.53855067 | 6.27E-13 | 0.43700006 | 9.10E-06 |
| Ccser1 | 0.50325306 | 1.16E-13 | 0.27665671 | 1.97E-08 |
| Grik1 | 0.47444385 | 2.33E-08 | 0.60762145 | 3.67E-12 |
| Fmnl2 | 0.4637757 | 1.06E-18 | 0.25127503 | 0.00113452 |
| Ap1s2 | 0.42845492 | 2.94E-12 | 0.2675064 | 1.39E-05 |
| Lrrc1 | 0.41017044 | 1.44E-11 | 0.32339497 | 2.18E-07 |
| Ptprk | 0.38407649 | 1.28E-11 | 0.33505535 | 3.74E-10 |
| Gabrb1 | 0.36728814 | 6.84E-10 | 0.36462969 | 6.27E-16 |
| Atp10a | 0.3587015 | 1.11E-07 | 0.29570341 | 1.76E-07 |
| Sub1 | 0.35534617 | 3.61E-07 | 0.26849379 | 6.04E-07 |
| Tmcc3 | 0.33516691 | 0.00011132 | 0.49566012 | 1.58E-19 |
| Arhgap6 | 0.31668323 | 1.66E-05 | 0.31039491 | 1.73E-09 |
| Arhgef28 | 0.28216655 | 0.00831711 | 0.39806128 | 8.27E-16 |
| Xist | 0.26640355 | 0.00014572 | 0.33486543 | 2.77E-11 |
| 4930555F03Rik | -0.2504493 | 0.00607082 | -0.3632657 | 7.63E-09 |
| Ntm | -0.2645728 | 2.40E-05 | -0.2803264 | 4.13E-05 |
| Sdc2 | -0.2981902 | 1.80E-08 | -0.3424908 | 5.11E-10 |
| Gm47283 | -0.3029916 | 8.22E-11 | -0.2519424 | 1.33E-06 |
| Ptprd | -0.3538745 | 3.68E-06 | -0.46183 | 2.28E-13 |
| Jun | -0.377936 | 0.00503601 | -0.4691269 | 2.93E-07 |
| Luzp2 | -0.4268187 | 2.43E-12 | -0.2525721 | 0.00093101 |
| Il1rapl1 | -0.4396021 | 4.11E-14 | -0.282823 | 9.24E-07 |
| Atp1a3 | -0.4516407 | 1.04E-20 | -0.3223472 | 6.53E-10 |
| Naaladl2 | -0.4667902 | 2.81E-18 | -0.26135 | 0.0002756 |
| Ldlrad4 | -0.5136721 | 5.78E-15 | -0.3398439 | 5.70E-06 |
| Fat3 | -0.5950315 | 3.65E-30 | -0.4087543 | 3.61E-14 |
| Tmx4 | -0.6519681 | 1.62E-29 | -0.4276142 | 1.26E-10 |

| gene | avg_log2FC_fasted | p_val_adj_fasted | avg_log2FC_lactating | p_val_adj_lactating |
|---|---|---|---|---|
| 6330403K07Rik | -0.686545 | 5.85E-44 | -0.2803992 | 4.29E-08 |
| Ncald | -0.7032342 | 1.80E-32 | -0.3455955 | 1.67E-08 |
| AC149090.1 | -0.7359445 | 1.32E-39 | -0.3793794 | 5.53E-08 |
| Tenm2 | -0.7532671 | 2.34E-25 | -0.4139553 | 3.81E-18 |
| Nsg2 | -0.8129794 | 6.76E-64 | -0.3689241 | 2.57E-14 |
| Lrrtm4 | -0.9695989 | 2.02E-48 | -0.252732 | 0.03680199 |
| Rbfox1 | -1.1959635 | 5.19E-58 | -0.5784536 | 2.04E-20 |
| Sgcz | -1.3503698 | 4.56E-87 | -0.4606414 | 1.58E-16 |
